# Mito-nuclear effects uncovered in admixed populations

**DOI:** 10.1101/349126

**Authors:** Arslan A. Zaidi, Kateryna D. Makova

**Affiliations:** Department of Biology, The Pennsylvania State University, University Park, PA 16802, USA

## Abstract

To function properly, mitochondria utilize products of 37 and >1,000 genes encoded by the mitochondrial and nuclear genomes, respectively, which should be compatible with each other. Discordance between mitochondrial and nuclear genetic ancestry could contribute to phenotypic variation in admixed populations. Here we explored potential mito-nuclear incompatibility in six admixed human populations from the Americas: African Americans, African Caribbeans, Colombians, Mexicans, Peruvians, and Puerto Ricans. For individuals in these populations, we determined nuclear genome proportions derived from Africans, Europeans, and Native Americans, the geographic origins of the mitochondrial DNA (mtDNA), as well as mtDNA copy number in lymphoblastoid cell lines. By comparing nuclear vs. mitochondrial ancestry in admixed populations, we show that, first, mtDNA copy number decreases with increasing discordance between nuclear and mitochondrial DNA ancestry, in agreement with mito-nuclear incompatibility. The direction of this effect is consistent across mtDNA haplogroups of different geographic origins. This observation suggests suboptimal regulation of mtDNA replication when its components are encoded by nuclear and mtDNA genes with different ancestry. Second, while most populations analyzed exhibit no such trend, in Puerto Ricans and African Americans we find a significant enrichment of ancestry at nuclear-encoded mitochondrial genes towards the source populations contributing the most prevalent mtDNA haplogroups (Native American and African, respectively). This likely reflects compensatory effects of selection in recovering mito-nuclear interactions optimized in the source populations. Our results provide the first evidence of mito-nuclear effects in human admixed populations and we discuss its implications for human health and disease.

## Introduction

Mitochondria participate in some of the most vital functions of eukaryotic cells, such as generation of ATP via oxidative phosphorylation (OXPHOS), regulation of calcium uptake, apoptosis, and metabolism of essential nutrients ^1^. Mitochondria harbor their own genome (mitochondrial DNA, or mtDNA) with 37 genes encoding 13 proteins essential for OXPHOS and the ribosomal and transfer RNAs required for their translation. Additionally, >1,000 nuclear genes encode proteins involved in mitochondrial function ^2,3^. The nuclear genome encodes most of the subunits of the OXPHOS complexes, and proteins required for replication and transcription of mtDNA. The nuclear-encoded mitochondrial genes must be transcribed in the nucleus, translated in the cytoplasm, and directed to mitochondria with the help of translocases and mitochondrial membrane proteins, which are themselves encoded by the nuclear genome ^2,4^. Thus, mitochondrial functions, and therefore many cellular functions in general, rely on fine-tuned interactions between mtDNA and the products of mtDNA-encoded and nuclear-encoded mitochondrial genes. As a result, we expect mtDNA and nuclear-encoded mitochondrial genes to coevolve, i.e. to undergo *mito-nuclear coevolution* ^5–11^. This expectation is especially plausible because of a smaller effective population size, higher mutation rate, and thus faster evolution, of mtDNA compared to the nuclear genome ^6^.

Evidence of mito-nuclear coevolution comes primarily from inter-population hybrids resulting from laboratory crosses of model organisms, such as fruit flies ^12–18^, marine copepods ^19–22^, and yeast ^23–25^. Inter-population hybrids in these organisms frequently exhibit reduced viability and fecundity ^16,17,19–21^. These phenotypes are associated with altered expression of OXPHOS genes ^13,26^, reduced OXPHOS activity, decreased ATP production ^14^, altered mtDNA copy number ^20,26^, and elevated oxidative damage ^21^. Fitness can often be restored if the hybrids are backcrossed with the maternal line but not with the paternal line ^20^, suggesting that their reduced fitness was caused by differences in ancestry between mitochondrial and nuclear genomes, hereafter called ‘m*ito-nuclear DNA discordance*’.

*Mito-nuclear incompatibility* –- which we define as any phenotypic manifestation of mito-nuclear discordance –- has also been observed in naturally occurring hybrids of diverging populations. For instance, Morales and colleagues (2016) analyzed two populations of the eastern yellow robin (*Eopsaltria australis*) –- which have adapted to different climates and carry different mtDNA haplotypes –- and discovered that highly differentiated regions of their nuclear genomes are enriched in nuclear-encoded mitochondrial genes. This observation suggests that divergence between the two populations has been maintained by mito-nuclear incompatibility, in spite of continued gene flow ^27^. Similarly, Baris and colleagues (2017) analyzed two killifish (*Fundulus heteroclitis*) populations with divergent mtDNA haplogroups, as well as hybrids between these two populations, and found that admixture fraction across differentiated nuclear loci in hybrids is associated with decreased OXPHOS activity ^28^. This finding points towards the role of mitochondrial and nuclear ancestry in altering the efficacy of mitochondrial function ^28^. Such studies demonstrate the utility of naturally occurring hybrid, i.e. admixed, populations in studying mito-nuclear coevolution and the phenotypic effects brought about by its disruption (i.e. by mito-nuclear incompatibility).

Currently, we know very little about the extent of mito-nuclear incompatibility and its contribution to phenotypic variation in humans. Several studies have shown that altered (due to mutations) interactions between mtDNA-and nuclear-encoded factors can modulate disease phenotypes for cardiomyopathy, predisposition to type 2 diabetes, as well as possibly for hearing loss and Huntington’s disease ^11,29^. Two additional studies have recently explored mito-nuclear incompatibility in humans in more detail. First, Sloan and colleagues (2015) tested for elevated mito-nuclear linkage disequilibrium (LD) across a set of 51 human populations from the Human Genome Diversity Project^30–32^. They hypothesized that as human populations diverged, certain allelic combinations between mtDNA and nuclear-encoded mitochondrial genes would have been disfavored if they were incompatible and if the effect sizes of such epistatic interactions were sufficiently large. However, they found no evidence of increased LD at single-nucleotide polymorphisms (SNPs) in such genes relative to the genomic background ^30^. In the second study, Sharbrough and colleagues (2017) investigated whether nuclear-encoded mitochondrial genes in modern humans are depleted in Neanderthal and Denisovan ancestry ^33^. Such a depletion is expected because previous studies have found no evidence of introgression of Neanderthal and Denisovan mtDNA into modern humans ^34,35^. If the divergence between humans and archaic hominins was sufficient to cause mito-nuclear incompatibility, selection could have purged archaic hominin alleles at nuclear-encoded mitochondrial genes because of incompatible or unfavorable interactions with human mtDNA. The authors found a significant underrepresentation of Neanderthal, but not Denisovan, ancestry at such genes in modern humans. These results suggest a complex history of mito-nuclear coevolution in modern humans and other hominins.

Studying recently admixed populations is another approach to test for mito-nuclear coevolution in humans. If mito-nuclear coevolution occurred in diverging human populations, then previously co-adapted mito-nuclear interactions could have been disrupted as a result of recent admixture, and mito-nuclear incompatibility might be observed in admixed populations at individual and population levels (Fig. 1). Within any given admixed individual (Fig. 1A), the nuclear genome, because of its biparental inheritance, recombination, and independent assortment, represents a mosaic of ancestry segments ^36,37^. However, mtDNA, because it is exclusively maternally inherited, maintains maternal ancestry only (Fig. 1A). Any discordance between mitochondrial and nuclear ancestry may cause mito-nuclear incompatibility within admixed individuals, depending on the degree of divergence between the ancestral populations and the extent to which the nuclear genome differs in ancestry from the mitochondrial genome (Fig. 1A).

**Figure 1.**
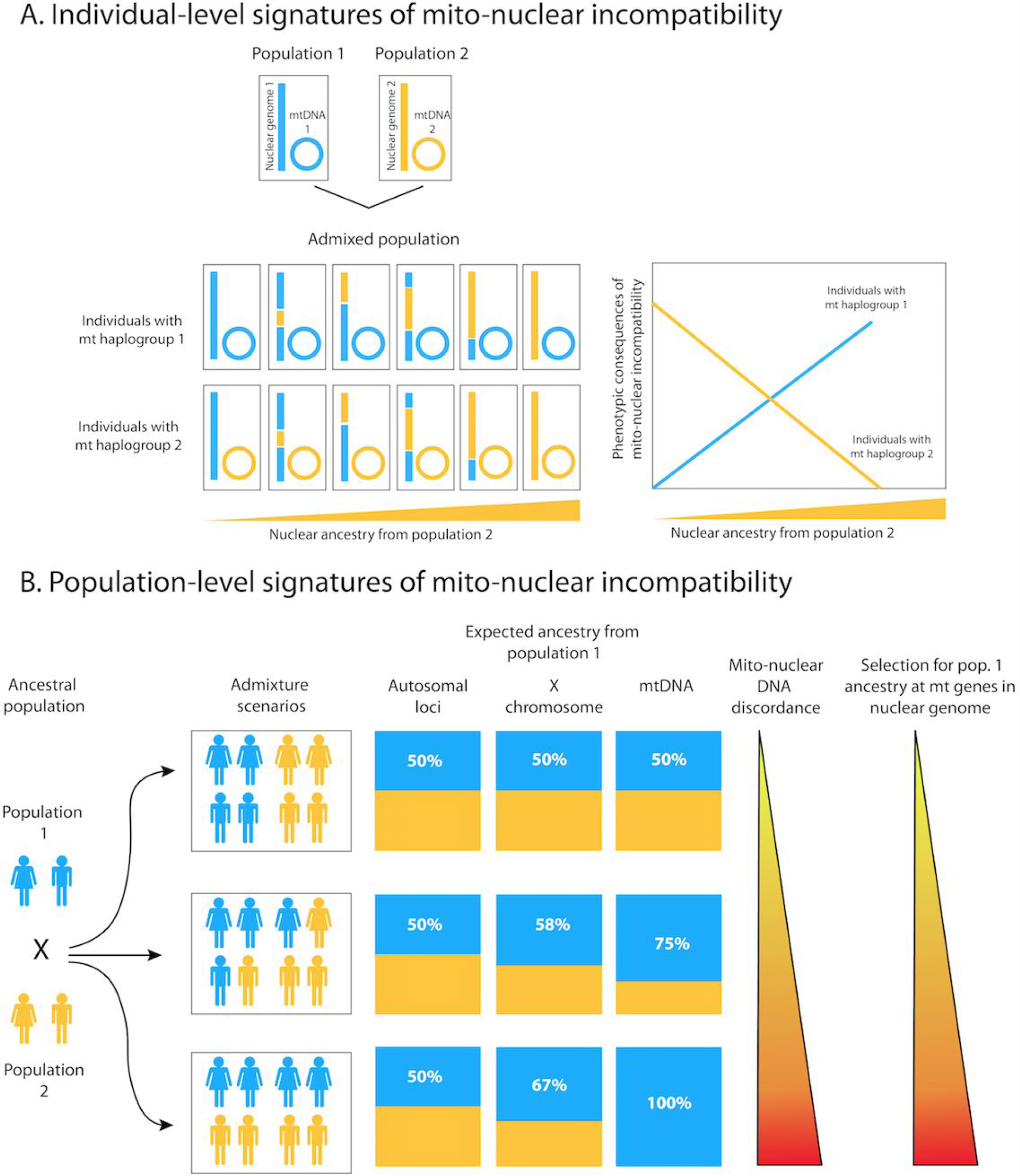
Expected signatures of mito-nuclear incompatibility at the level of (A) individuals or (B) populations. **(A)** The nuclear genome of admixed individuals might contain varying levels of ancestry from multiple ancestral populations, whereas mtDNA derives its ancestry from only one population. Increasing discordance (concordance) between mitochondrial and nuclear ancestry in individuals can lead to increasing (decreasing) levels of mito--nuclear incompatibility, which can have phenotypic consequences. **(B)** Sex--biased admixture can lead to varying levels of ancestry proportions on different chromosome types based on their modes of inheritance. In extreme cases, the mtDNA ancestry can be entirely from one population even though the nuclear ancestry is highly admixed. This discordance in ancestry between the nuclear and mitochondrial genomes might result in selective pressure for ‘matching’ ancestry at nuclear--encoded mitochondrial genes.

Signatures of mito-nuclear coevolution can also manifest at a population level (Fig. 1B), particularly if there is sex bias in the genetic contributions from the source populations, as in the case of admixture in the Americas. Colonization and slave trade in the last 500 years resulted in admixture among Native Americans, Africans, and Europeans in the Americas. Historical and genetic studies agree that the early European settlers were primarily males. This population history, compounded with the social stratification that resulted in directionally skewed gene flow between European males and African and Native American females, has resulted in differences in the frequency of European ancestry among the autosome, X chromosome, Y chromosome, and mtDNA ^38–47^. Increasing levels of sex-biased admixture can lead to increasing discordance between mitochondrial and nuclear ancestry proportions in admixed populations (Fig. 1B). If coadapted mito-nuclear combinations are disrupted in admixed populations, leading to a reduction in fitness, then we expect selection to act towards restoring them, i.e. to shift the average ancestry fraction at nuclear-encoded mitochondrial genes in favor of the source population contributing the highest proportion of females (Fig. 1B).

In this manuscript, we explored signatures of mito-nuclear incompatibility and coevolution in admixed human populations from the Americas using the data from the 1,000 Genomes Project (1000 Genomes Project Consortium et al. 2015). We first tested whether the discordance between mitochondrial and nuclear ancestry in an individual’s genome has an effect on their mtDNA copy number, a cellular phenotype that is a known biomarker for many health-related outcomes, including aging, fertility, and several types of cancers ^48–51^. Next, we investigated whether within admixed populations there are systematic shifts in ancestry frequency at nuclear-encoded mitochondrial genes towards the source population contributing the highest proportion of mtDNA. Our results present evidence of mito-nuclear incompatibility, and suggest presence of selection geared towards overcoming it, in human admixed populations. Thus, our observations support the notion that mito-nuclear coevolution has occurred in both non-admixed and admixed human populations.

## Materials and Methods

### mtDNA copy number estimation

Given the average sequencing depth of the autosomes and mtDNA, and the fact that there are two autosomal chromosome copies per cell, we can compute the number of copies of mtDNA per cell:

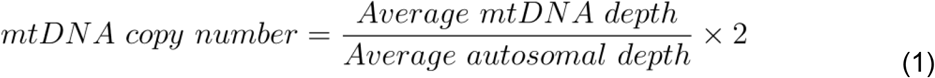

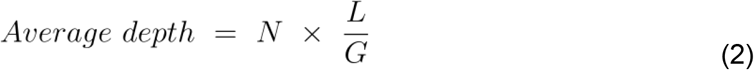

In the equation for average depth, *N* is the total number of reads aligning to the chromosome, *L* is the average length of a read, and *G* is the size of the chromosome in base pairs. We first calculated mtDNA copy number for each autosomal chromosome separately, and then calculated the mean across all chromosomes.

A subset of samples (a total of 24) in the 1000 Genomes Project Data were sequenced at both low (2-4x) and high coverage (20-40x). We used these to validate whether mtDNA copy numbers calculated from the low-vs. high-coverage alignments agree with each other. As shown in Fig. S1, the copy numbers generally agree, with a few exceptions. Some samples show appreciable mtDNA copy numbers when calculated using the high-coverage alignments but low copy numbers when calculated using the low-coverage alignments (Fig. S1). The source annotation of these samples indicates that some of them (samples for which this information is available) were sequenced from peripheral blood mononuclear cells (PBMCs), instead of Lymphoblastoid Cell Lines (LCLs) (Fig. S1, Table S1). The difference in mtDNA copy number seen between the two cell lines is consistent with previous observations that LCLs are known to carry significantly higher mtDNA copy numbers than PBMCs ^52–54^. Because annotation for the source DNA is not available for all samples (Table S1), we plotted the density of mtDNA copy number calculated from the low-coverage alignments and observed a clear separation between samples sequenced from PBMCs and LCLs (Fig. S2). We removed all samples with less than 250 mtDNA copies per cell from the downstream analysis to exclude samples which were sequenced from PBMCs in an effort to limit variation due to DNA source. After removing such samples, the correlation coefficient between mtDNA copy number from low-coverage and high-coverage alignments is 0.71, as opposed to 0.66 before removing them.

#### Global ancestry and mtDNA haplogroup

We downloaded the 1000 Genomes phase 3 vcf files and retained individuals who belonged to the ACB, ASW, CEU, CLM, MXL, PEL, PUR, or YRI populations in subsequent analyses. Global ancestry was calculated using ADMIXTURE ^55^. For this purpose, we merged the 1000 genomes genotype data with the genotype data from Native American groups published by Mao and colleagues ^56^. We included SNPs that overlapped across both datasets (a total of 691,435 SNPs) and converted the genotypes to binary format for use with Plink ^57,58^. We further removed palindromic (A/T, G/C) SNPs to ensure strand consistency across both datasets. Subsequently, the two datasets were merged and SNPs were pruned for LD (r^2^ threshold of 0.1), which resulted in 88,442 SNPs. We ran unsupervised ADMIXTURE ^55^ analysis on this genotype dataset for k = 1, 2, 3, 4, and 5 and used the ancestry proportions for k = 3 for all downstream analyses, since it had the lowest cross-validation error (Fig. S3). The three ancestry components, C1-3, correspond to European, African, and Native American ancestry, respectively (Fig. 2A-B).

**Figure 2.**
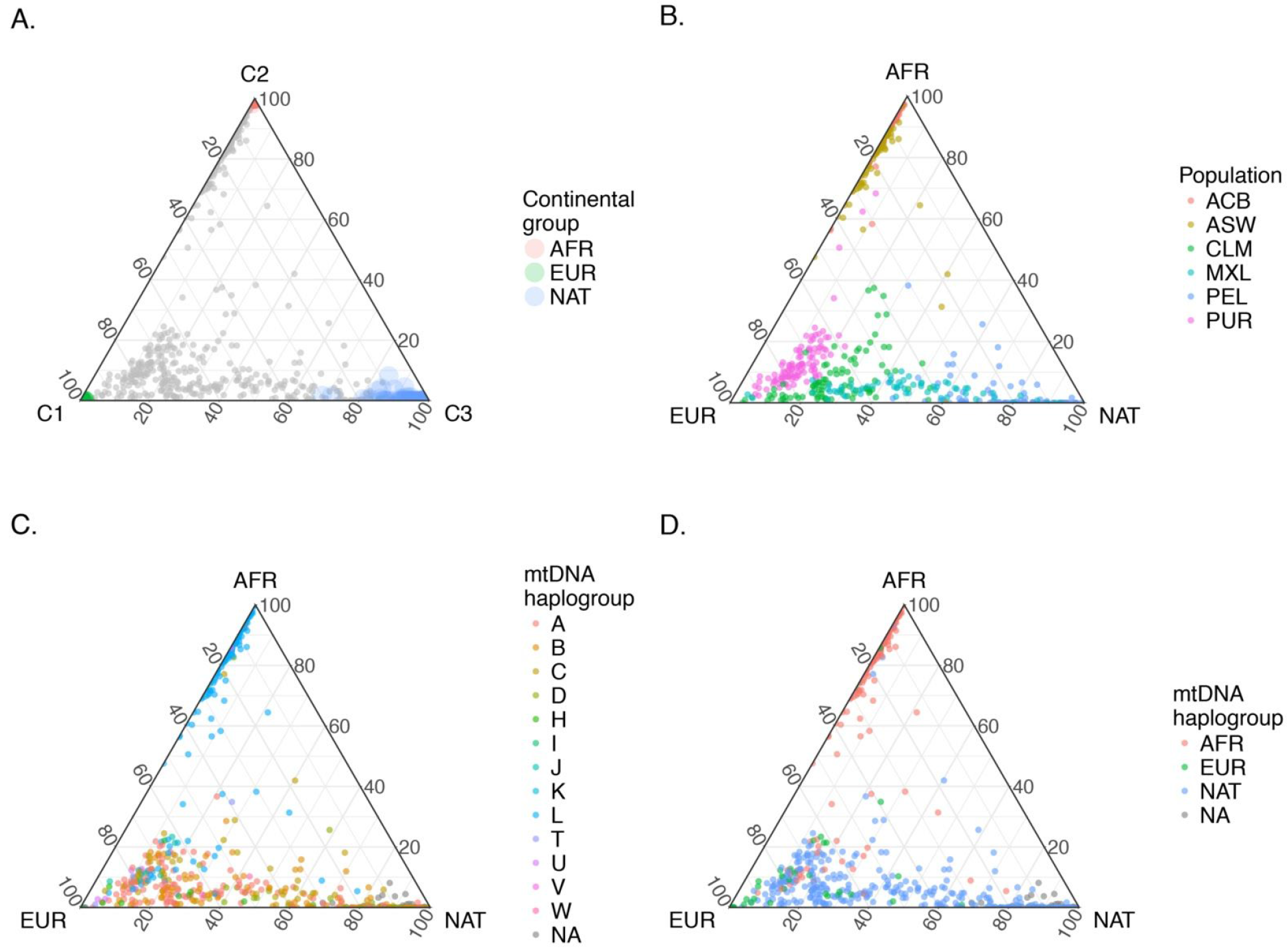
Ternary plots showing the distribution of African, European, and Native American ancestry in the samples analyzed. **(A)** ADMIXTURE components 1, 2, and 3 correspond to European, Native American, and African ancestry, respectively. The grey points are the admixed samples from the 1,000 Genomes dataset and the colored points are samples serving as proxies for source populations. **(B)** The ancestry structure of admixed populations. **(C)** Distribution of mtDNA haplogroups among admixed individuals. **(D)** mtDNA haplogroups grouped by region where they are thought to have been most commonly found prior to admixture. NAT –- Native American, other abbreviations are spelled out in Table 1.

**Table 1.** The number of individuals from each population used in the study. NA - data not available or not used in this study.

| Ancestry group | Population | Number of individuals with sequence alignments | Number of individuals with genotype data | Data Source |
| --- | --- | --- | --- | --- |
| Admixed | African Americans from the Southwest (ASW) | 60 | 61 | 1,000 Genomes Project Data<sup>68</sup> |
|  | African Caribbean from Barbados (ACB) | 96 | 96 | 1,000 Genomes Project Data<sup>68</sup> |
|  | Mexicans from Los Angeles (MXL) | 67 | 64 | 1,000 Genomes Project Data<sup>68</sup> |
|  | Peruvians (PEL) | 86 | 85 | 1,000 Genomes Project Data<sup>68</sup> |
|  | Puerto Ricans (PUR) | 105 | 104 | 1,000 Genomes Project Data<sup>68</sup> |
|  | Colombians (CLM) | 94 | 94 | 1,000 Genomes Project Data<sup>68</sup> |
| African | Yoruba from Nigeria (YRI) | 109 | 108 | 1,000 Genomes Project Data<sup>68</sup> |
| European | Utah residents of Northern and Western European ancestry (CEU) | 99 | 99 | 1,000 Genomes Project Data<sup>68</sup> |
| Native American | Aymara | NA | 25 | Mao et al.<sup>56</sup> |
|  | Nahuan | NA | 14 | Mao et al.<sup>56</sup> |
|  | Quechua | NA | 24 | Mao et al.<sup>56</sup> |
|  | Maya | NA | 25 | Mao et al.<sup>56</sup> |
| Total |  | 717 | 799 |  |

We used Haplogrep version 2.02 ^59^ to determine the mtDNA haplogroup, for individuals from the 1000 Genomes Project only, as the individuals from Mao et al. ^56^ were not genotyped for mitochondrial variants. All mtDNA haplogroups were called with high accuracy (minimum posterior probability of 0.78). To increase statistical power, we grouped together haplogroups belonging to the same major haplogroup (e.g. L1b1a3 was grouped with L3d1b1 under the L major haplogroup; Fig. 2C). We further grouped major haplogroups into regional groups, corresponding to pre-colonization origins (L: African; A, B, C, D: Native American; H, J, K, T, U, V, W: European) (Figs. 2C-D, S4). We excluded two individuals (HG01272 and NA19982), whose mtDNA was predicted to belong to the M haplogroup most frequently found in South Asia. This way, both nuclear and mitochondrial ancestry was categorized into only three regional groups: Native American, European, and African (Fig. 2D).

#### Local ancestry

Local ancestry for autosomes was generated using RFMix ^60^ by the 1000 Genomes Project admixture working group as described by Martin and colelagues ^61^ https://personal.broadinstitute.org/armartin/tgp_admixture/snp_pos/). For downstream analyses, we masked out regions of the genome where local ancestry was called with less than 0.9 maximum posterior probability.

#### Sex-biased admixture

The relative contribution of males and females from each of the three relevant ancestral groups (African, European, and Native American) was inferred by comparing the ancestry fractions estimated from the X chromosome and the autosome using the approach described in ^62^. If 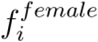 is the proportion of ancestors of the admixed group, who were female and from population *i*, and 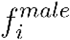 who were male and from population *i*, we assume that for each admixed group (e.g. PUR, CLM etc.), 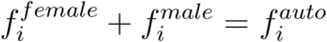, the mean autosomal ancestry fraction from population *i*, where *i* ε {African, Native American, European}. Furthermore, we assume that 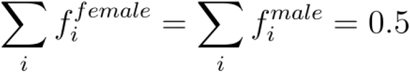. Thus, for values of 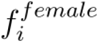 and 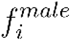, the expected ancestry fraction for the X chromosome in the population, 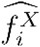 is ^63^:

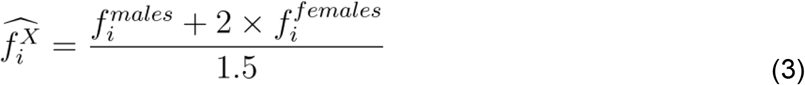

We performed a grid search for the values of 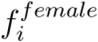 and 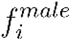 that equal 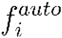 and minimize the squared deviation between 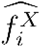, predicted using equation (3), and the mean ploidy-adjusted X-chromosomal ancestry fraction inferred from genotype data. Estimated values of 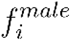 and 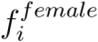 and confidence intervals around these estimates, generated by bootstrapping, are shown in Fig. S5.

#### Simulation of drift since admixture

We simulated the expected amount of drift in local ancestry since admixture in Puerto Ricans and Colombians using a simple hybrid-isolation demographic model ^64,65^ where all three ancestry groups mix at some time in the past in proportions equal to the mean global ancestry averaged across individuals in the population. The resulting population is then allowed to mate randomly for *g* generations, simulated by drawing 2*N* autosomes, and *N/2* copies of mtDNA, in each generation with the probability of drawing a locus of European, African, or Native American ancestry determined by the relative ancestry proportions in the previous generation. After *g* generations, the final ancestry frequency at each locus, averaged across individuals in the population, is recorded. We repeated this process 10,000 times to simulate the amount of drift for 10,000 independent loci. We used 17 generations for *g* in PUR and 14 generations for *g* in CLM, similar to values estimated for these populations by Gravel et al. (2013), and 1,250 for *N* following Tang et al. (2007). While our model is simple and does not reflect the true demographic history of the populations, which likely involved continued gene flow for many generations as well as complex, non-random mating patterns, our goal is not to infer the true demographic history of these populations but solely to simulate the amount of drift in local ancestry since admixture. As we show in Fig. S6, the simulated data match the observed distribution of local ancestry quite well.

#### Local ancestry enrichment in nuclear-encoded mitochondrial genes

First, we calculated the frequency of Native American, European, and African ancestry at every SNP in the genome by averaging across all individuals in a population. These were subtracted from the the mean ancestry fraction across all SNPs to calculate the deviation in local ancestry at each SNP. We used the list of genes from MitoCarta 2.0 and split the list into mitochondrial and non-mitochondrial genes according to the classification provided ^66^. We further split the mitochondrial genes into two subsets to separately analyze a list of 167 genes curated by Sloan et al. (2015), which code for proteins that are part of the replication and transcription machinery of mtDNA, and the ribosomal and OXPHOS complexes in the mitochondria. We classified this list of genes as ‘High-mt’ and the remaining mitochondrial genes as ‘Low-mt’. An unweighted block bootstrap approach was used to generate the distribution of mean deviation in local ancestry for each gene category. We generated windows of 5 Mb spanning each gene (± 2.5 Mb on either side of a gene’s midpoint) to take into account LD among SNPs. Subsequently, we used bedtools ^67^ to intersect SNPs, at which local ancestry deviation was calculated previously, with these windows. For each gene category, 167 windows were sampled with replacement, to match the number of genes in the smallest category (i.e. High-mt), and the mean ancestry deviation was calculated, first for each window, and then across windows. This process was repeated 1,000 times to generate a distribution of mean deviation in local ancestry for each gene category.

## Results

### Nuclear and mitochondrial ancestry proportions in the admixed populations

To study mito-nuclear incompatibility and coevolution in humans, we analyzed genomic alignments and genotype data from six admixed populations that are part of the 1000 Genomes Project ^68^: (1) African Americans from the Southwest (ASW); (2) African Caribbeans from Barbados (ACB); (3) Colombians from Medellin, Colombia (CLM); (4) Mexicans from Los Angeles (MXL); (5) Peruvians from Lima, Peru (PEL); and (6) Puerto Ricans from Puerto Rico (PUR). We also used data from Utah residents of Northern and Western European ancestry (CEU) and from Yorubans from Ibadan, Nigeria (YRI), who serve as proxies for the European and African source populations, respectively. To represent the Native American ancestry component, we analyzed previously published genotype data ^56^ from the following four groups: (1) Aymara; (2) Nahuan; (3) Quechua; and (4) Maya. The data set is summarized in Table 1.

We determined global ancestry –- the overall contribution of African, European, and Native American ancestry to the nuclear genome –- of each analyzed individual using ADMIXTURE (Fig. 2A-B; see Methods for details). In agreement with previous studies ^44,61^, the individuals from admixed populations derive their genetic ancestry from three primary source populations: Native American, European, and African (ADMIXTURE cross-validation error is lowest at *k* = 3; Fig. S3; see Methods). The proportion of ancestry from each source population varies among the admixed populations (Fig. 2B; Table S2 and S3) because of differences in admixture histories ^44,45^. We also determined the mtDNA haplogroup for each individual (Table S4; see Methods for details). Among the admixed individuals, African (L) and Native American (A, B, C, and D) mtDNA haplogroups are more frequent than European haplogroups (H, J, K, T, U, V, and W; Fig. S4), consistent with female bias in the non-European contribution ^69^.

### mtDNA copy number decreases with increasing mito-nuclear DNA discordance

We hypothesized that increasing discordance between nuclear and mitochondrial ancestry in admixed individuals will lead to an increase in the degree of incompatibility (Fig. 1), for instance, between mtDNA origins of replication and nuclear-encoded mtDNA replication machinery ^70,71^ and therefore, to a decrease in mtDNA replication efficiency. If our hypothesis is correct, then mtDNA copy number should decrease with increasing degree of mito-nuclear DNA discordance (Fig. 1A). To evaluate this prediction, we determined the mtDNA copy number from sequence alignments of DNA extracted from lymphoblastoid cell lines (LCLs; see Methods). We computed mtDNA copy number for each individual from the six admixed populations as well as from the CEU and YRI populations (which were used as proxies of the European and African source populations, respectively (there are no ‘non-admixed’ Native Americans who are part of the 1000 Genomes Project Dataset ^68^). An advantage of using LCLs to study mtDNA copy number variation is that they exhibit high mtDNA content and elevated expression of genes involved in mtDNA replication and transcription, as well as of respiratory genes, consistent with elevated mitochondrial biogenesis ^72^. Furthermore, because they are maintained following standard protocols in a laboratory, variation due to differences in individuals’ environments, from whom the LCLs are derived, is unlikely to systematically confound our analysis of mtDNA copy number.

We found that mtDNA copy number decreases as nuclear ancestry becomes increasingly dissimilar to mtDNA ancestry (Fig. 3A and Fig. 4), consistent with our hypothesis (Fig. 1A). To obtain this result, we regressed mtDNA copy number against the degree of discordance between mtDNA ancestry and nuclear ancestry, as measured by the fraction of the nuclear genome that is from a different geographical origin than the mtDNA haplogroup. For instance, for individuals with Native American mtDNA haplogroups, mito-nuclear DNA discordance is the proportion of nuclear ancestry that is not Native American (i.e. African plus European). There is a significant negative correlation between mtDNA copy number and mito-nuclear DNA discordance in admixed individuals (Fig. 3A; *Beta* = −0.193, *one-sided P-value* = 1.14 × 10^−04^, *r* = −0.19). The intercept of this slope, i.e. when mito-nuclear DNA discordance is zero, is similar to the median mtDNA copy number in the individuals from source populations, CEU and YRI, who are not admixed (Fig. 3B). The negative correlation between mtDNA copy number and mito-nuclear DNA discordance in admixed individuals is consistent across mtDNA haplogroups from three different geographic origins –- Native American (*Beta* = −1.06; *P* = 1.53 × 10^−05^), African (*Beta* = −0.63; *P* = 0.026), and European (*Beta* = −0.30; *P* = 0.370) –- even though it is not always statistically significant. Moreover, in each case, copy number for mtDNA of one geographic origin decreases with increasing nuclear ancestry from each of the other two geographic origins (panels outside of the top-left to bottom-right diagonal in Fig. 4). For instance, for individuals with Native American mtDNA haplogroups, the mtDNA copy number decreases with increase in both African and European ancestry. Conversely, as the ancestry between mtDNA and the nuclear genome becomes more similar, mtDNA copy number increases (top-left to bottom-right diagonal in Fig. 4). The lack of power, especially in admixed individuals carrying European haplogroups, is likely due to the limited female European contribution (Fig. 4). Overall, our results show that a phenotype, mtDNA copy number, is negatively correlated with mito-nuclear DNA discordance, consistent with mito-nuclear incompatibility in admixed individuals (Fig. 1A).

**Figure 3.**
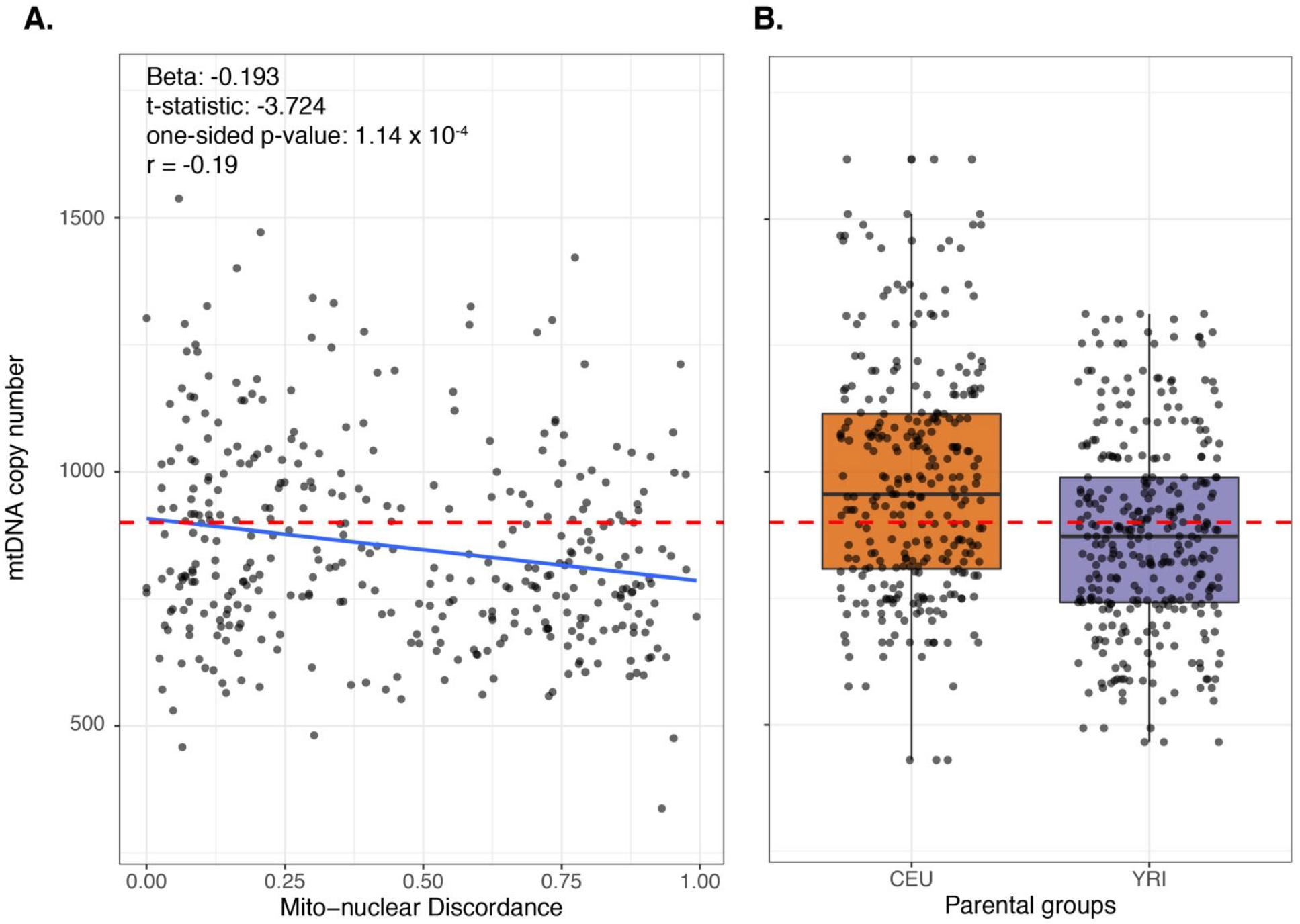
mtDNA copy number in (A) admixed and (B) non-admixed populations. **(A)** mtDNA copy number is negatively correlated with the discordance between mitochondrial and nuclear DNA ancestry. Standardized beta coefficient, t-statistic, one-sided P-value, and correlation coefficient are shown. The discordance score is one minus the ancestry proportion from the same source population as for the mtDNA haplogroup. The red dashed line is the median mtDNA copy number calculated across the individuals from CEU and YRI source populations, plotted separately in **(B)**.

**Figure 4.**
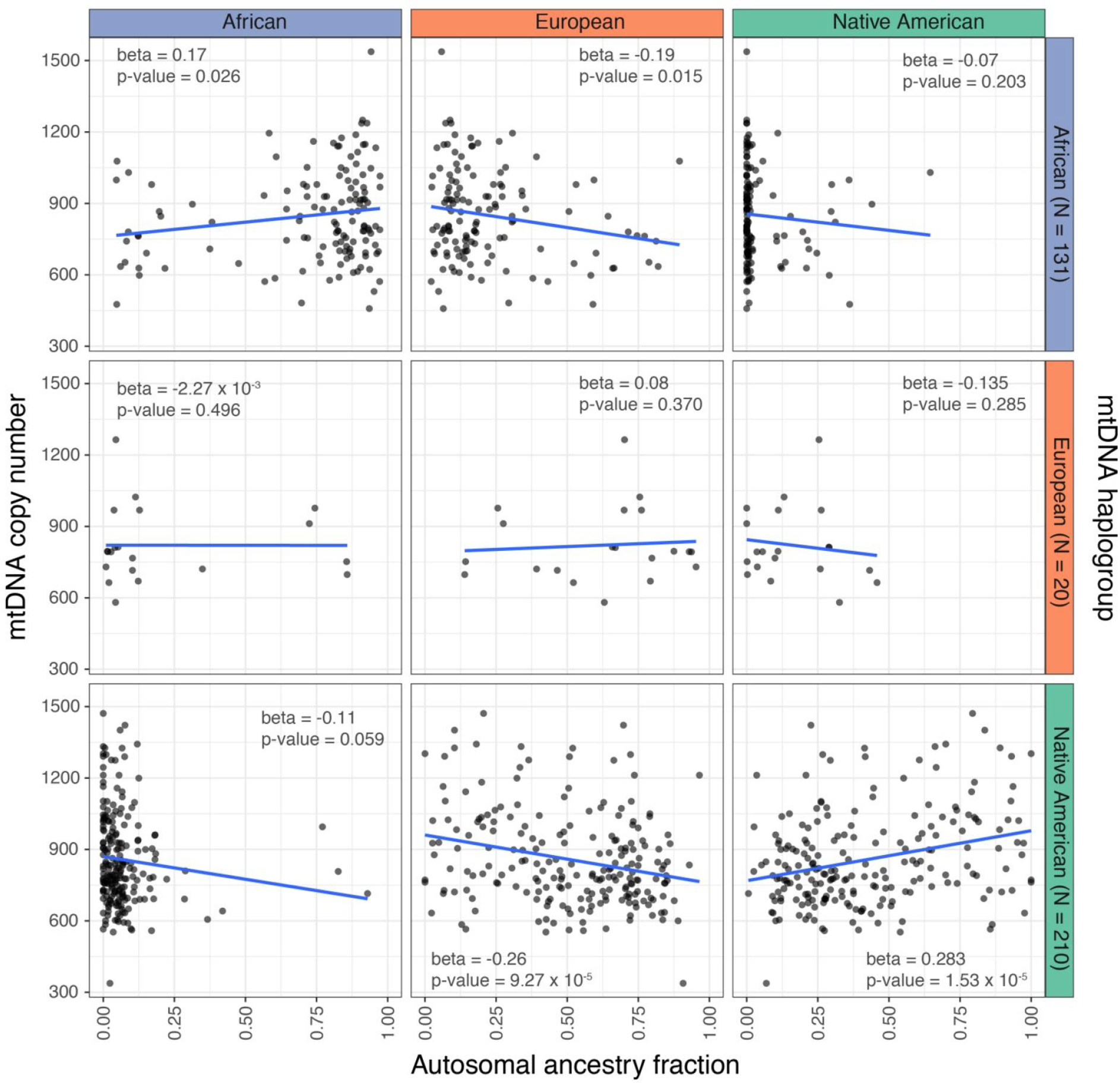
Scatter plots of mtDNA copy number (y--axis) against autosomal ancestry fraction (x--axis) Rows and columns are split by the geographic origin of the mtDNA haplogroup and source populations, respectively. The top- left to bottom -right diagonal, where the nuclear and mtDNA ancestry are similar, shows positive correlations between ancestry and mtDNA copy number, while the other panels show negative correlations, consistent with our predictions in Fig. 1A. Beta coefficients and two-sided P-values are shown.

### Sex-bias in admixture inferred from the nuclear and mitochondrial genomes

According to our second hypothesis, we expect mito-nuclear DNA discordance at a population level to increase with increasing degree of sex bias in admixture (Fig. 1B). For example, if population 1 contributes only females and population 2 contributes only males to the admixed population (third admixture scenario in Fig. 1B), the expected percentage of mtDNA ancestry from population 1 is 100% compared to 50% for autosomal loci. Our results corroborate previous studies ^44–46,61^, suggesting that gene flow in the Americas was sex-biased. To quantify the degree of sex bias from a source population, we estimated the male and female contributions to each admixed group from that source population, by comparing the global ancestry proportions on the autosomes and the X chromosome (Figs. 5 and S5; see Methods). In all of the six admixed populations, the European contribution is male-biased and non-European (African and Native American) contribution is female-biased. For example, there were likely more males, than females, who contributed European ancestry to Peruvians (Fig. 5).

**Figure 5.**
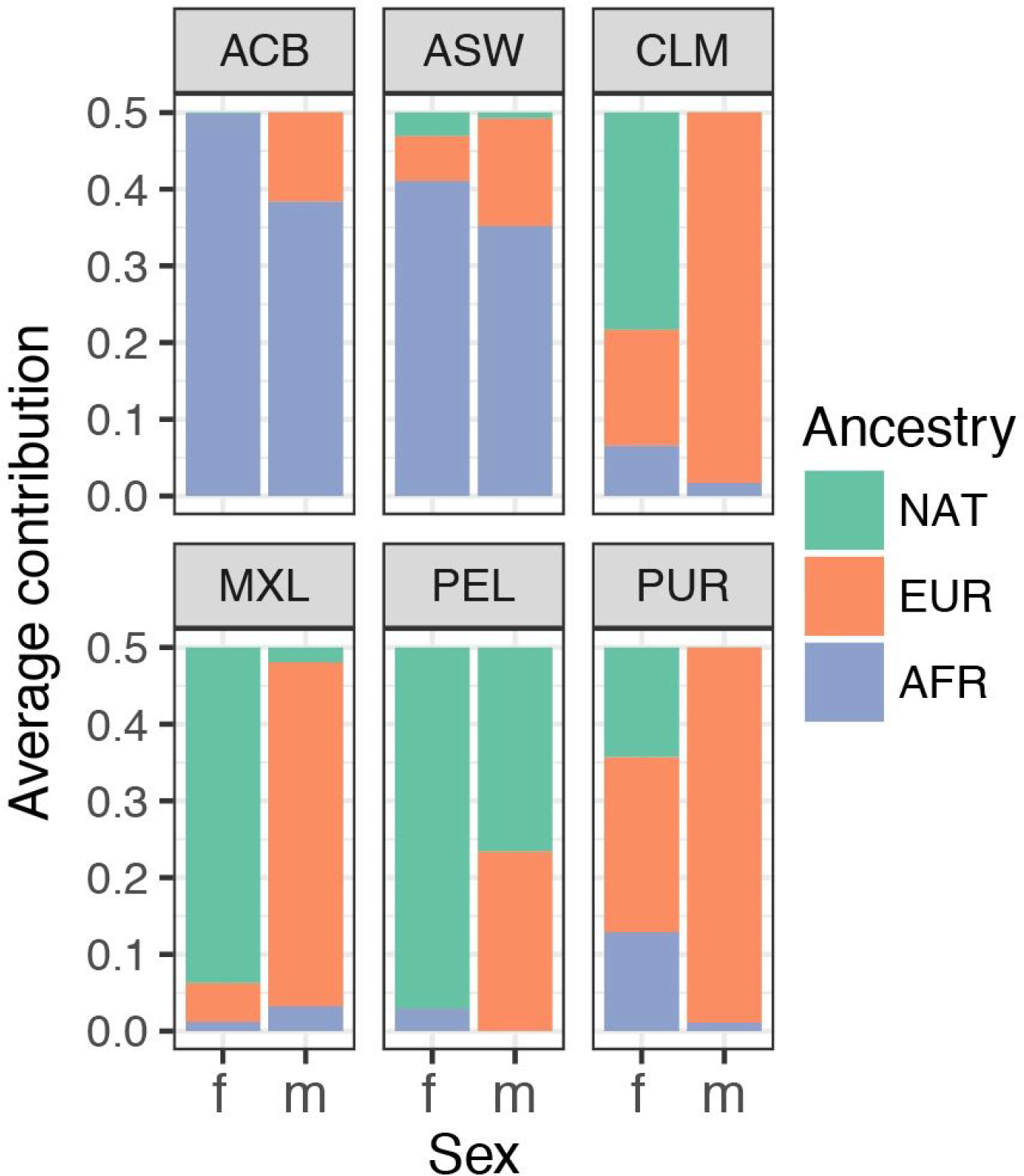
Estimated female and male composition in the ancestral populations for each admixed group.

We further compared the observed proportion of mtDNA haplogroups of African, European, or Native American origin with the estimated proportion of contributing females from these source populations as inferred from the nuclear genome (Fig. 6). Both independently measure the female contributions and thus are expected to be similar. We found that, in most admixed populations analyzed, the observed frequency of mtDNA haplogroups falls within the expected distribution (generated using a non-parametric bootstrap, see Methods) of the proportion of females from each source population (Fig. 6). However, in Colombians (CLM) and Puerto Ricans (PUR), the frequencies of Native American mtDNA haplogroups are much higher and, concomitantly, the frequency of European haplogroups are much lower, than expected (Fig. 6). At least two factors can explain this result. First, since mtDNA represents a single genealogical history, it yields a ‘noisier’ estimate of the proportion of females from each parental population than the estimate based on autosomal and X-chromosomal loci, which represent multiple genealogical histories because of recombination and independent assortment. Second, we expect larger fluctuations in mtDNA ancestry as a result of drift because of its smaller effective population size compared to that of autosomal or X-chromosomal loci. To test whether genetic drift since admixture can account for the deviation in mtDNA frequency in Puerto Ricans and Colombians, we simulated the amount of drift expected for mtDNA in these populations based on their admixture history. In both Colombians and Puerto Ricans, drift since admixture is not sufficient to account for the observed deviation in mtDNA ancestry (Fig. S7).

**Figure 6.**
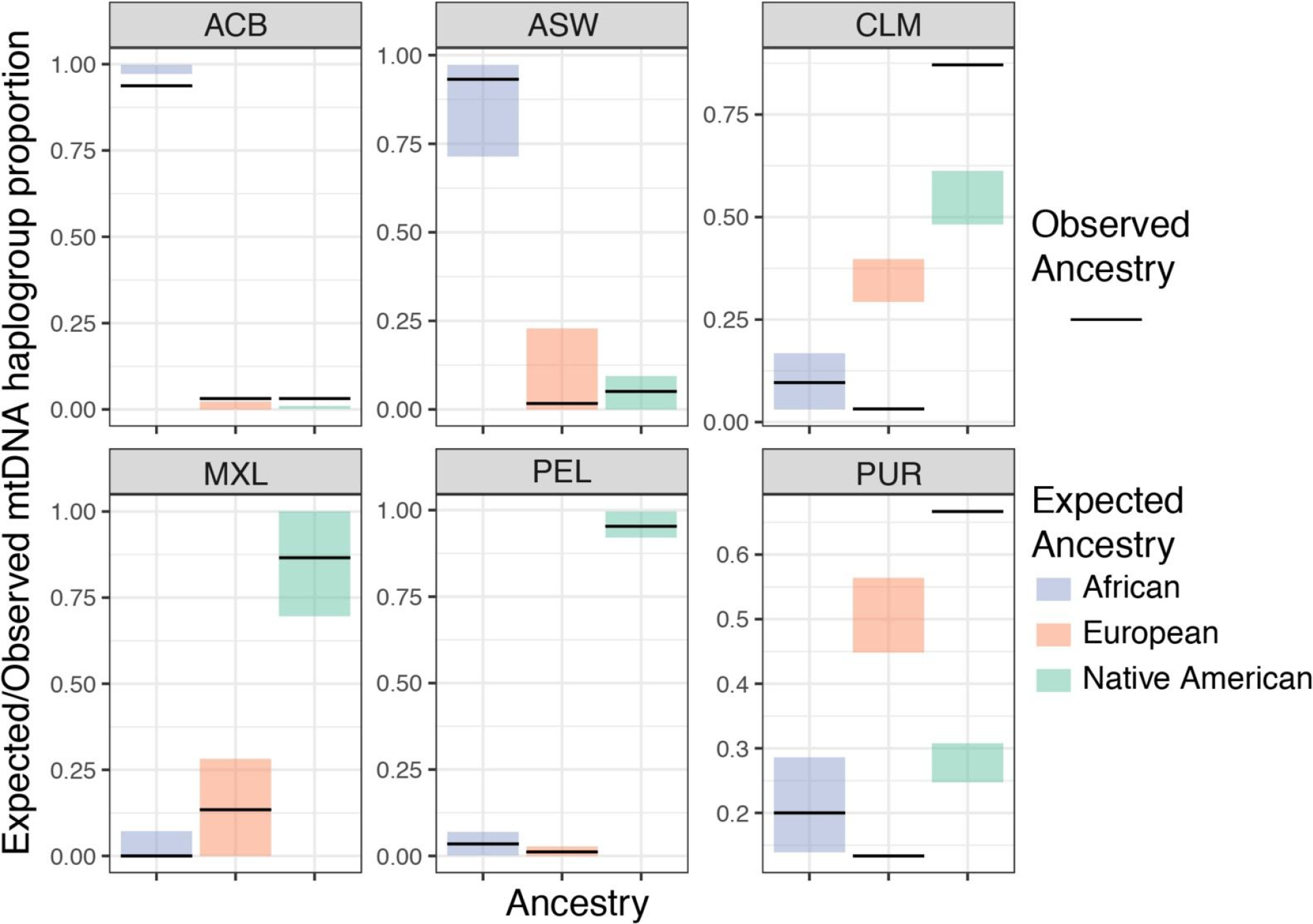
Expected vs. observed frequency of mtDNA haplogroups from each of the three source populations. The colored boxes show the 95% bootstrap interval of the expected mtDNA haplogroup frequency based on the proportion of ancestral females estimated from the ancestry proportions on the X- chromosome and autosomes (Fig. 5). The black horizontal lines show the observed frequency of mtDNA haplogroup. While in most cases the observed mtDNA haplogroup frequency falls within range of expectations, in some cases (i.e. in CLM and PUR), the observed frequencies deviate greatly from the expectation.

### Selection on local ancestry at nuclear-encoded mitochondrial genes

Based on our second hypothesis, we expected selection at nuclear-encoded mitochondrial genes to favor ancestry from the source population contributing the highest proportion of females, to compensate for potentially maladaptive mito-nuclear combinations resulting from recent admixture. For example, in Puerto Ricans, since there is a high frequency of Native American mtDNA, we expect selection to favor Native American ancestry at nuclear-encoded mitochondrial genes. Our ability to detect such a signature at individual loci is limited because of: (1) the large number of genes involved in mitochondrial function (>1000) ^2^, (2) the low amount of genetic differentiation among human populations (Fst ≈ 0.1) ^73^, (3) only a relatively recent admixture history in the populations analyzed ^44,45,61^, and (4) the relatively small sample size of our data set. However, we might be able to detect systematic shifts in the ancestry frequency for nuclear-encoded mitochondrial genes as a group. To test for such a signal of ancestry enrichment at nuclear-encoded mitochondrial loci, for each admixed population analyzed, we first calculated the deviation in local ancestry at each SNP by subtracting the global Native American, European, and African ancestry proportion in that population (see Methods). Subsequently, we downloaded a list of nuclear genes from MitoCarta 2.0 ^66,74^ and split them into mitochondrial (N = 960) vs. non-mitochondrial (N = 17,456) based on their classification (Table S5). The mitochondrial genes encode proteins with experimental evidence of mitochondrial localization, whereas the non-mitochondrial genes have no such evidence ^66^. We next followed a published approach ^30^ and further split the 960 mitochondrial genes into two subsets –- 167 high-confidence, or ‘High-mt’, genes (genes encoding proteins that are part of the mtDNA replication and transcription machinery, and of ribosomal and OXPHOS complexes) and remaining 793 ‘Low-mt’ genes (Table S5). We used a block bootstrap approach to generate a distribution of the mean deviation in local ancestry for each functional gene category (for 167 High-mt, 793 Low-mt, and 17,456 non-mitochondrial genes, see Methods for more details). Based on our hypothesis, local ancestry should deviate from neutral expectations in favor of the source population contributing the highest proportion of mtDNA (Fig. 1B).

We find that in three out of six admixed populations analyzed, the mean ancestry of nuclear-encoded mitochondrial genes does not significantly deviate from expectation (the 95% bootstrapped confidence interval spans the zero line; Fig. 7), consistent with no evidence of selection for local ancestry at such genes. However, we found a significant enrichment in Native American ancestry at High-mt genes in Puerto Ricans, and an enrichment in African ancestry at High-mt genes in African Americans (Fig. 7). Because Native American mtDNA haplogroups are more frequent in Puerto Ricans, and African mtDNA haplogroups are more frequent in African Americans (Fig. 5), these results are consistent with the predictions of our hypothesis (Fig. 1B). However, we also observe a significant enrichment in European ancestry at High-mt genes in Mexicans (Fig. 7), which contradicts our hypothesis. This lack of a consistent pattern of enrichment across admixed populations paints a complex picture of the selection regimes acting on nuclear-encoded mitochondrial genes. It should be noted that the systematic shifts in ancestry that we observe for the High-mt genes are inconsistent with sampling artifacts because we do not observe such patterns for non-mitochondrial genes (category ‘Non-mt’ in Fig. 7).

**Figure 7.**
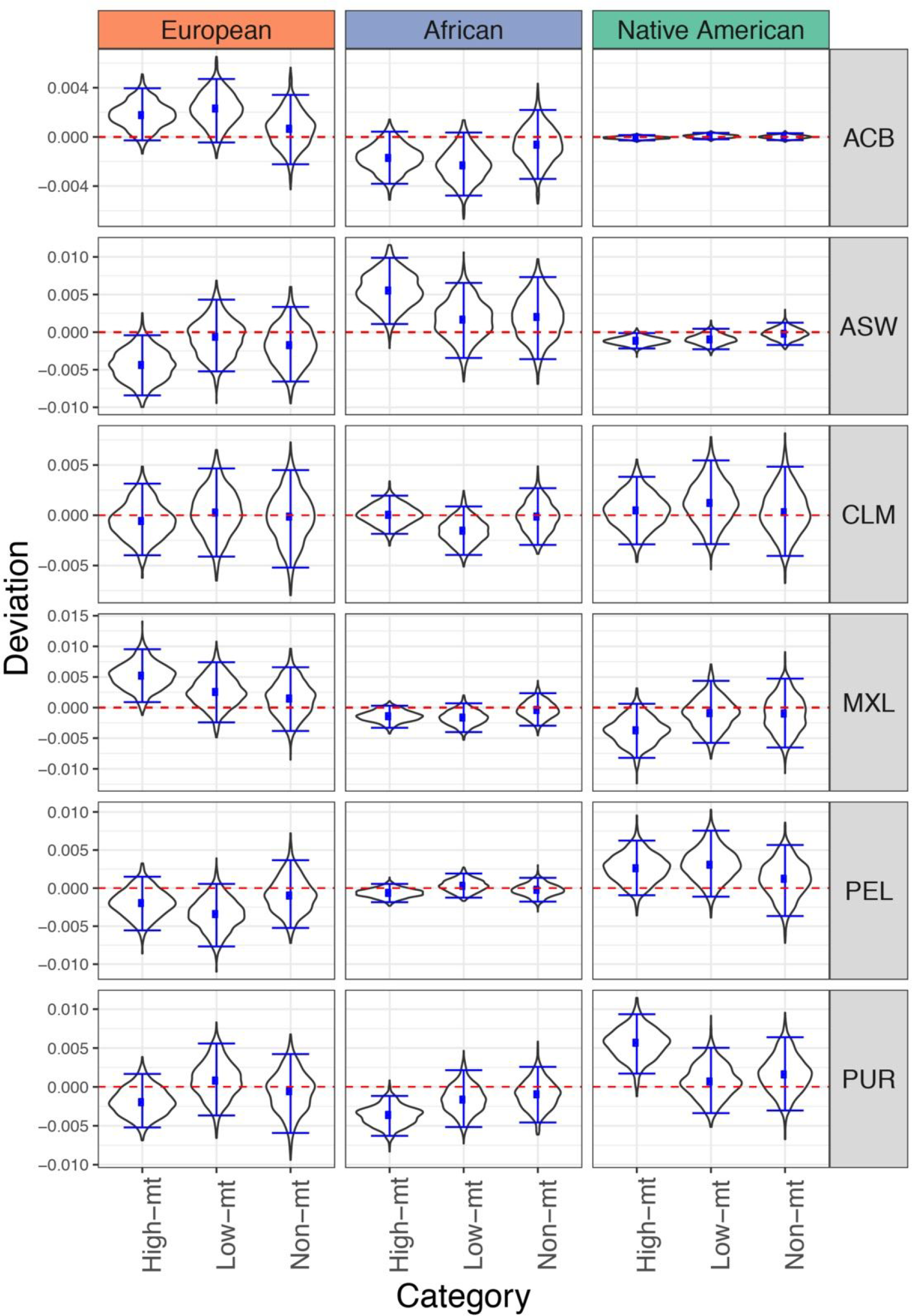
Systematic deviations in local ancestry for different functional categories of genes. The y--axis shows local ancestry deviation and the x--axis lists the functional categories. ‘High-mt’ (167 genes) are nuclear genes that encode important subunits of mitochondrial replication, transcription and the OXPHOS complexes. ‘Low-mt’ (793 genes) are nuclear genes that were inferred to have mitochondrial function by Mitocarta 2.0 ^66^, but are not part of the ‘High-mt’ gene set. ‘Non-mt’ (17,456 genes) are genes that do not have known or inferred mitochondrial function based on Mitocarta 2.0 ^66^. A block bootstrap approach (see Methods) was used to generate the distributions. Briefly, we sampled 167 windows of 5 Mb spanning the genes, with replacement, within each category and calculated the mean local ancestry deviation across these windows. This was repeated for 1,000 bootstraps to generate the distributions.

## Discussion

The potential contribution of epistatic interactions between nuclear- and mtDNA-encoded mitochondrial genes to phenotypic variation in human populations is poorly understood. Admixed populations provide a unique opportunity to investigate this question. Because of the difference in the mode of inheritance between mitochondrial and nuclear DNA, and the frequently sex-biased nature of admixture, the ancestry between mitochondrial and nuclear genomes is frequently mismatched in admixed individuals. Such a discordance can potentially disrupt fine-tuned interactions between mtDNA-encoded and nuclear-encoded mitochondrial genes. We investigated the potential consequences of the mito-nuclear DNA discordance in six admixed American populations represented in the 1,000 Genomes Project. Due to complex admixture history ^44,45,61^, the degree of discordance in ancestry between mtDNA and nuclear DNA differs both across populations and among individuals from the same population.

First, we tested for effects of mito-nuclear DNA discordance on the phenotype of admixed individuals. We found that mtDNA copy number, a biomarker for several phenotypes ^48^, decreases as the nuclear and mtDNA become increasingly dissimilar in ancestry. Because mito-nuclear compatibility is important for regulation of mtDNA copy number, its reduction due to mito-nuclear DNA discordance might reflect the incompatibility between the origins of mtDNA replication and nuclear-encoded proteins involved in mtDNA replication ^75^. Interestingly, several of the studied haplogroups –- B, D, H, T, U and J –- have fixed differences in the experimentally established mtDNA recognition sites for proteins of the replication machinery ^76^. These substitutions could lead to population differences in mtDNA binding affinity of polymerase-**γ** (POLG), mtDNA helicase (TWINKLE), and mitochondrial single strand binding proteins (mtSSB), three of the major proteins involved in mtDNA replication ^71^, which themselves also exhibit substantial sequence diversity in humans (unpublished data from the Makova Lab). Whether this diversity in the mtDNA recognition sites and in the replication machinery is the result of mito-nuclear co-evolution remains to be tested. Future functional experiments should validate effects of mtDNA haplogroup and nuclear DNA combinations on mtDNA copy number directly. This can be performed, for instance, in cybrids –- cellular hybrid lines carrying the same nuclear genetic background but different mtDNA haplotypes ^77^. Cybrids carrying different mito-nuclear ancestry combinations would also be helpful in elucidating whether mito-nuclear DNA discordance leads to higher mutation and heteroplasmy burden in mtDNA. Nevertheless, we have shown that mito-nuclear DNA discordance contributes to phenotypic variation in admixed individuals, despite relatively low differentiation among human populations ^73,78^.

Even though a number of biological and technical factors can influence mtDNA copy number, they are unlikely to bias our results in a systematic way. Mitochondrial biogenesis and decay are dynamic processes which can change in response to environmental factors, such as nutrient availability, oxidative stress, and temperature ^79–81^. While such factors can covary with ancestry in admixed populations (e.g. socioeconomic status, a predictor of stress, is highly correlated with ancestry in Latino populations ^82^), we do not think that they have significantly affected our results. This is because we measure mtDNA copy number in lymphoblastoid cell lines (derived from peripheral blood cells) that have been cultured under standard laboratory conditions for long periods of time. Thus, environmental variation among cells is minimal and does not reflect the initial variability that existed among samples when they were first collected. Additionally, even though many environmental variables can covary with ancestry in admixed populations, they could not have confounded our results as the ancestry proportions of the cells were unknown when they were first cultured. Thus, biological variation, minimal due to standardized cell culturing, and technical variation, due to sampling and sequencing, would add some noise to the measurement of mtDNA content rather than a systematic bias and would not lead to the observed correlation between mito-nuclear DNA discordance and mtDNA copy number, a pattern consistent across mtDNA haplogroups. This correlation should be replicated and explored in future studies, ideally with mtDNA copy number measured across biological and technical replicates to reduce noise. It would also be interesting to test the effect of mito-nuclear DNA discordance on other mitochondrial phenotypes such as mitochondrial morphology and rate of ATP production.

Next, we leveraged sex-biased admixture to study effects of mito-nuclear incompatibility at the population level. Consistent with previous studies ^44–46,61^, we found a significant sex bias in admixture for all six admixed populations studied. In particular, more European males than females, and more African and/or Native American females than males contributed to admixture. In most admixed populations we found a high level of concordance between the female contribution estimated from nuclear genetic markers and that estimated from mtDNA, for each source population. However, in Colombians and Puerto Ricans, the proportion of Native American ancestry in mtDNA is significantly higher than expected based on the ancestry information in the nuclear genome. This deviation cannot be explained by genetic drift experienced by mtDNA since admixture due to its smaller effective population size, and could be due to selection for the Native American haplogroups in Colombians and Puerto Ricans, if the Native American mtDNA was better adapted to the environment. Indeed, climate adaptation may have played a role in mtDNA diversification across human populations ^83,84^. Differences in the local environment among admixed populations might explain why we do not observe similar deviations in the other admixed populations.

The assumed model of admixture dynamics has several limitations, which could lead to biased estimates of the female and male contributions. Specifically, we assume a hybrid-isolation model with equal reproductive variance and similar generation times between males and females. First, models that incorporate continuous gene flow are likely to yield slightly different estimates, especially if admixture occurred recently, i.e. within the last five generations (Goldberg et al. 2015). Because admixture for the populations used in this study started much earlier (>10 generations ago, ^44,45,61^), this is not a major concern in our case. Second, shorter generation times in females than in males would result in a smaller effective population size for the mtDNA compared to the nuclear genome ^85^. This would lead to stronger drift in mtDNA, which might explain the observed discrepancy in ancestry proportions between mtDNA and the nuclear genome in Colombians and Puerto Ricans without invoking selection. Third, men typically tend to have higher variance in reproductive success relative to women ^86,87^, which would increase the effective population size of the mtDNA compared to the nuclear genome and might render the observed discrepancies in Colombians and Puerto Ricans non-significant. A detailed discourse of how these competing processes affect the inference of sex bias in admixture dynamics is beyond the scope of this paper, but needs to be addressed in future studies. Despite these concerns, it is clear that the frequency of non-European mtDNA haplogroups is much higher than the frequency of non-European ancestry at nuclear loci, in all six populations.

Finally, we hypothesized that the discordance in the ancestry frequency between mtDNA and nuclear loci would lead to selection on nuclear-encoded mitochondrial genes due to mito-nuclear coevolution. In three out of the six admixed populations studied (African Caribbeans, Colombians, and Peruvians, we found the ancestry of nuclear-encoded mitochondrial genes to be consistent with the expectation based on female contribution estimated from mtDNA data (Fig. 7). This could mean that there has been no selection on mito-nuclear interactions in these populations or that we do not have power to detect it because of drift and/or relatively limited sample size. We observed an enrichment of Native American ancestry in Puerto Ricans, and an enrichment of African ancestry in African Americans, at nuclear-encoded mitochondrial genes. Because Puerto Ricans predominantly carry Native American mtDNA haplogroups, whereas African Americans primarily carry African haplogroups, this observation is consistent with our expectation of selection against mito-nuclear incompatibility in admixed populations. However, we also found a significant enrichment of European ancestry in Mexicans –- another population with predominantly Native American mtDNA haplogroups. This observation is inconsistent with selection against mito-nuclear incompatibility in admixed populations. This complex picture among different admixed populations suggests that other competing selective pressures on mitochondrial genes might be involved and it may not be straightforward to detect signatures of mito-nuclear incompatibility from deviations in local ancestry alone.

Our ability to detect mito-nuclear incompatibility signatures is influenced by several factors including degree of sex bias, time since admixture, effective population size, selection strength, and the number of loci under selection. One potential limitation of this analysis is that we assume that all nuclear-encoded mitochondrial genes have equal effect sizes for mitochondrial function. Since mitochondrial function is a highly complex trait, this assumption is likely incorrect. A more accurate way of testing for selection at these genes would be to weigh the contribution of each locus by its effect size. Unfortunately, we do not know what these effect sizes are because genome-wide association studies of mitochondrial phenotypes, such as mtDNA copy number and rate of ATP production, are yet to be conducted in humans. While systematic collection of such data for large cohorts of individuals is pending, it would also be highly informative to explore the effects of mito-nuclear interactions on various health-related phenotypes in large-scale datasets such as the UK Biobank ^88^.

In conclusion, our results demonstrate that discordance between mtDNA and nuclear ancestry for mitochondrial genes can affect the phenotype of admixed individuals. Note, however, that the observed effect on mtDNA copy number is relatively small and needs to be explored in other cells and tissue types. Other phenotypes potentially affected by mito-nuclear DNA discordance, e.g. ATP production, should be examined as well. Mito-nuclear DNA discordance contributes to disease phenotypes of non-admixed individuals (reviewed in ^11^, therefore we expect this phenomenon to contribute even more to disease variation in admixed individuals, and this needs to be evaluated in future studies. Such evaluation is also critical for making advances in mitochondrial replacement therapy (MRT), a technique in which the mtDNA carrying disease-associated mutations in a patient’s oocyte is replaced with mtDNA from a healthy donor oocyte ^89–91^. Despite the success of MRT, many human and non-human primate embryos created via mitochondrial replacement do not develop normally ^89^. Mitochondrial replacement can also lead to detrimental effects on growth, development, respiration, metabolism, aging, fertility, and survival in non-primate animals ^89^. Human cybrid lines also show variation in mtDNA copy number, ATP turnover rates, reactive oxygen species production, and expression of OXPHOS genes ^92^. Despite these observations, the degree to which the mitochondrial haplogroup of a donor should match the genetic background of the ‘nuclear’ parents in MRT in humans remains unanswered. The answer to this question is even less clear for admixed individuals whose nuclear genomes are a mix of ancestry from different populations. Our work highlights the potential of studying admixed individuals to better understand phenotypic effects of mito-nuclear DNA discordance, which will be useful in elucidating MRT-associated risks and evaluating disease susceptibility in contemporary admixed and non-admixed populations ^11,93^.

## Description of Supplemental Data

Supplementary data include eight figures and five tables.

## Declaration of Interests

The authors declare no competing interests.

## Acknowledgments

We thank Rasmus Nielsen, Mark Shriver, and William Chase for their comments on the manuscript. This project was supported by a seed grant awarded to AAZ and KDM from the Center of Human Evolution and Development (CHED) at The Pennsylvania State University, and by a grant from NIH (R01GM116044). Additional funding was provided by Penn State Eberly College of Sciences, The Huck Institute of Life Sciences at Penn State, and the Penn State Institute for CyberScience, as well as, in part, under grants from the Pennsylvania Department of Health using Tobacco Settlement and CURE Funds. The department specifically disclaims any responsibility for any analyses, responsibility, or conclusions.

## Supplementary Figures

**Figure S1.**
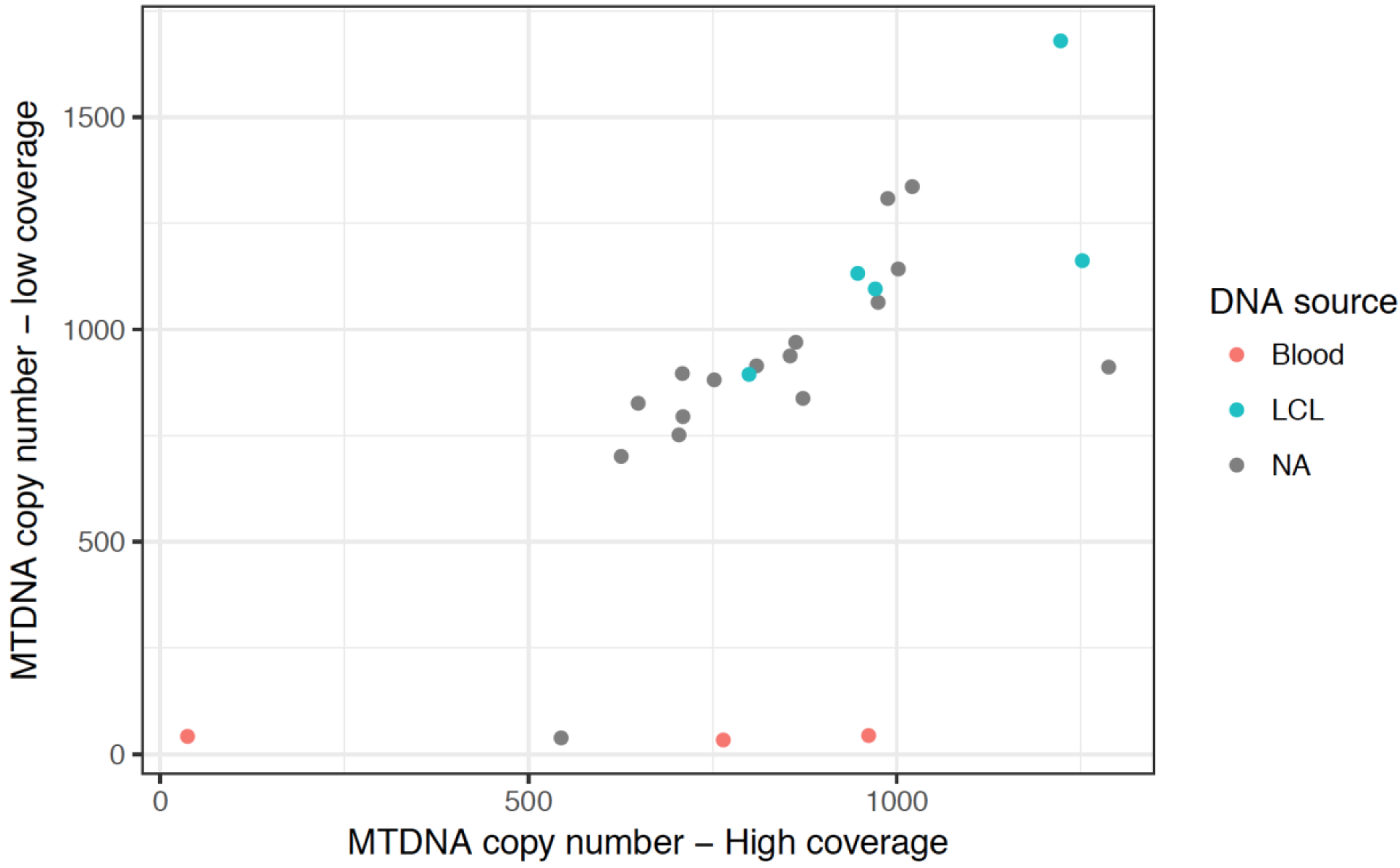
Plot showing concordance between mtDNA copy number calculated from high-coverage and low-coverage alignments. The source of the DNA sample (where available) is indicated. LCL - lymphoblastoid cell line

**Figure S2.**
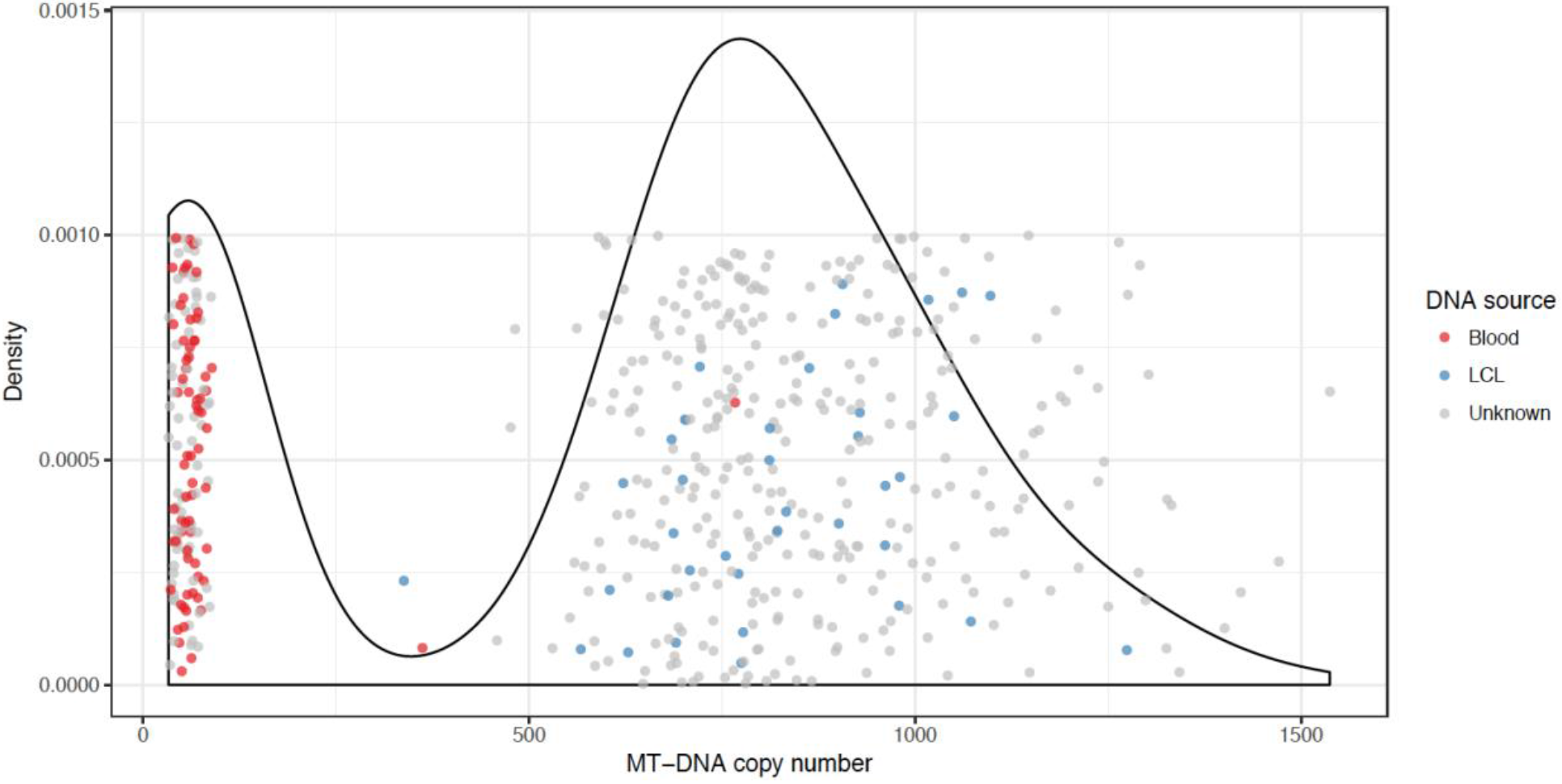
Distribution of mtDNA copy number calculated using low-coverage sequence alignments. There is a clear separation between samples sequenced from lymphoblastoid cell lines (LCLs) and peripheral blood mononuclear cells (PBMCs). In two cases, samples from lymphoblastoid cells appear to be mislabeled as blood.

**Figure S3.**
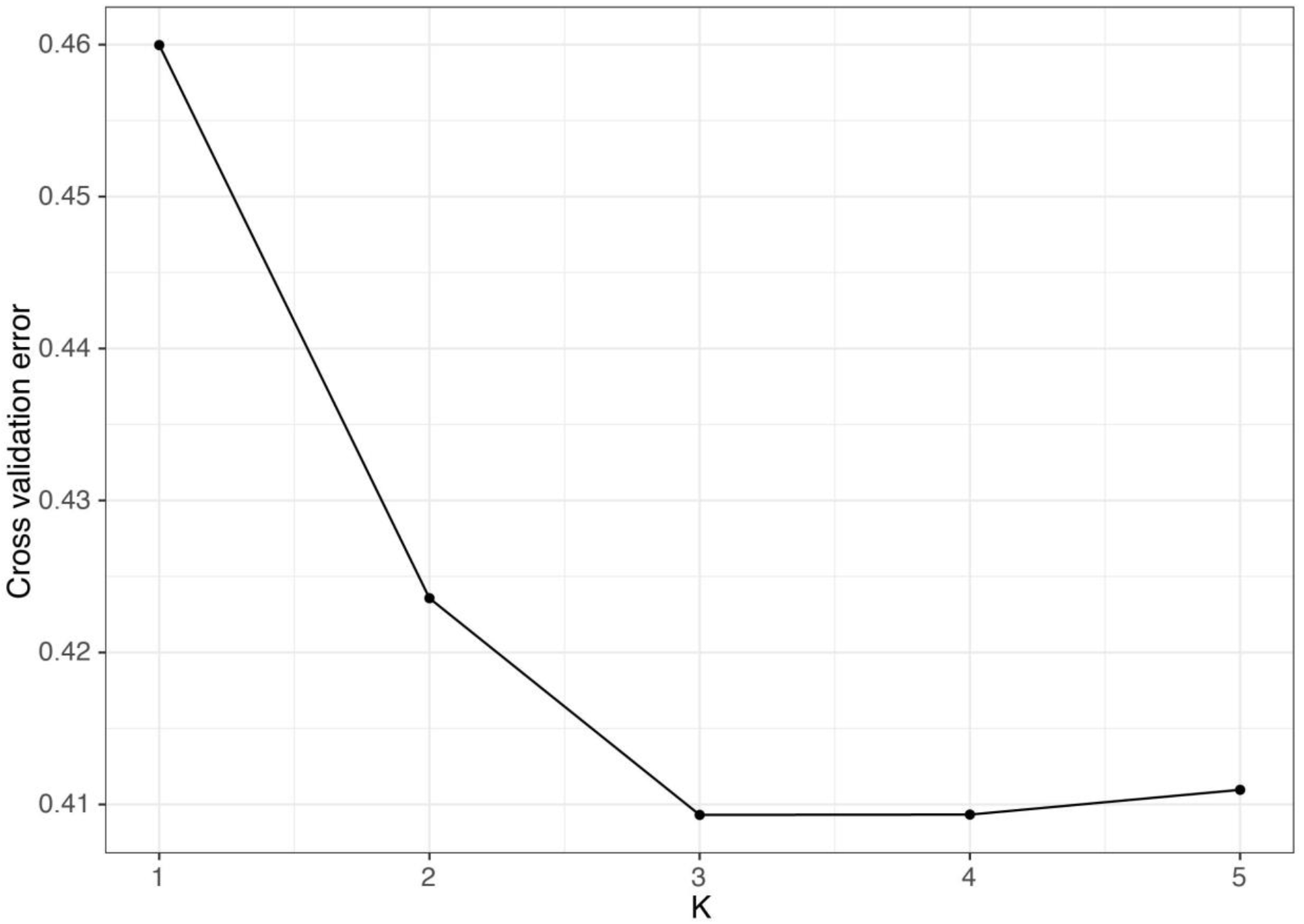
ADMIXTURE cross-validation error for values of K from 1 to 5. Cross-validation error is lowest for k = 3.

**Figure S4.**
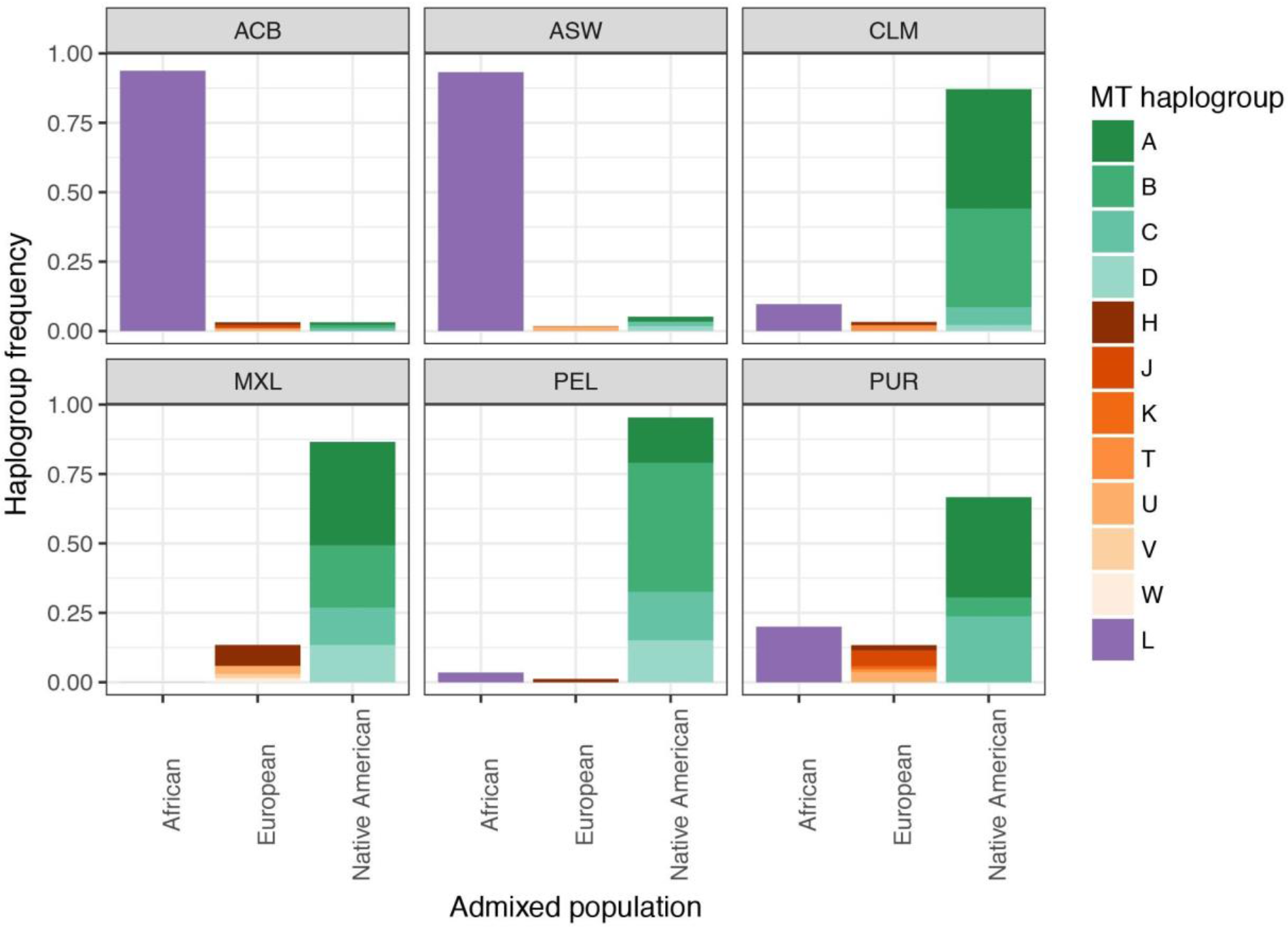
Breakdown of mtDNA haplogroup frequency observed in each admixed pop

**Figure S5.**
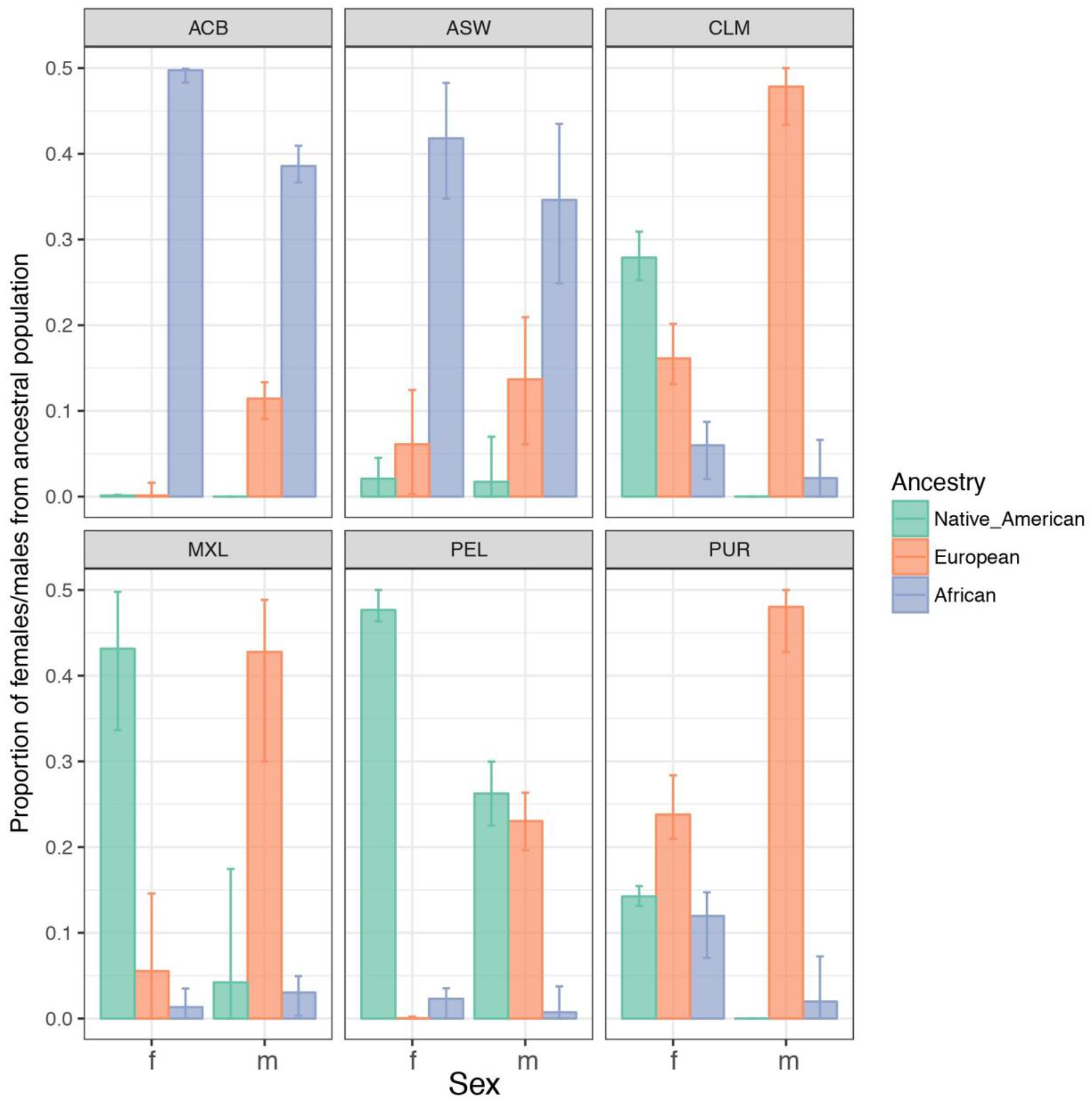
Estimates of male and female contributions from source populations with bootstrapped confidence intervals.

**Figure S6.**
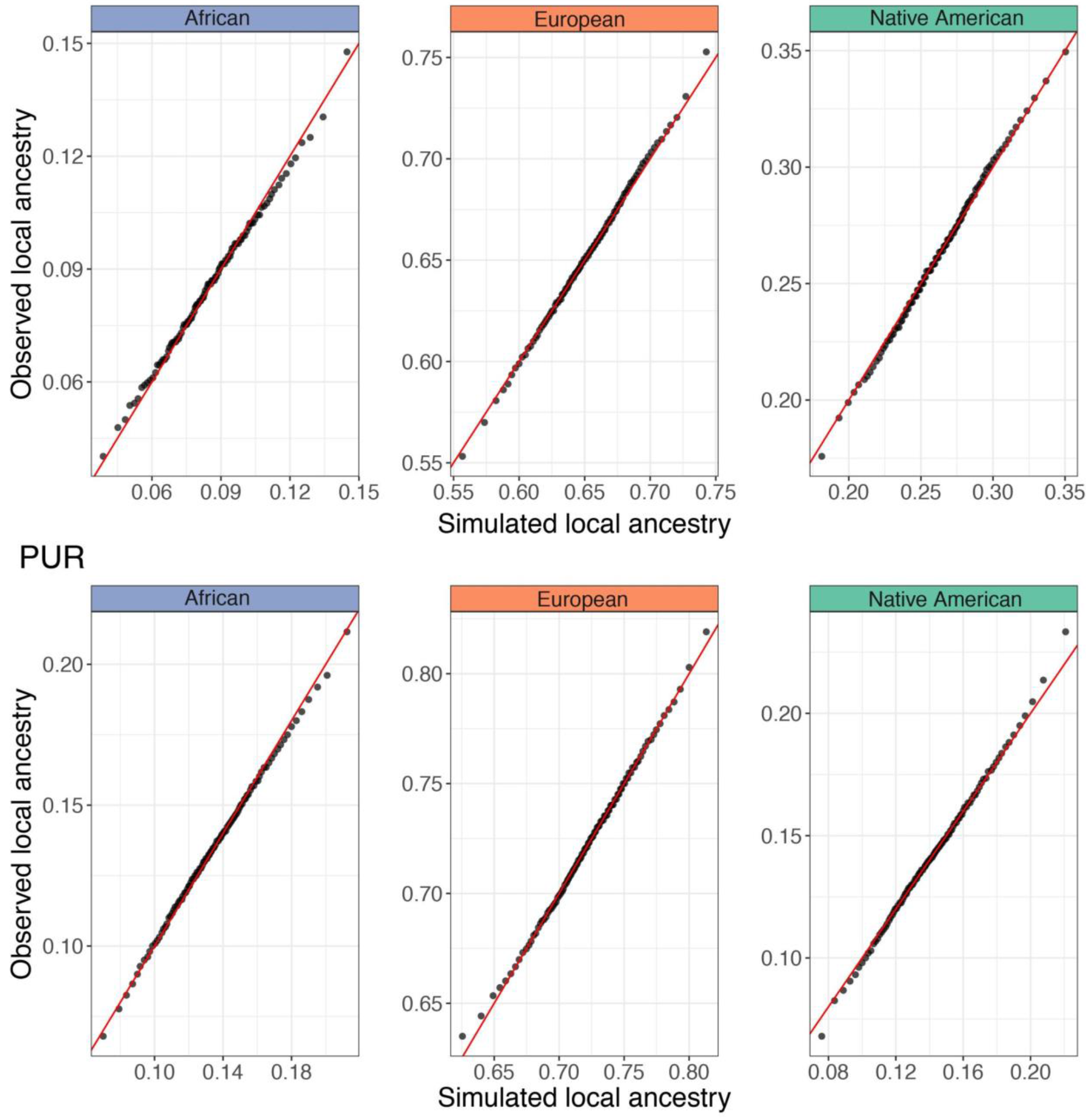
The simulated amount of drift in local ancestry (for 1,000 independent loci) is similar to the observed amount of drift in local ancestry at autosomal loci in both PUR and CLM.

**Figure S7.**
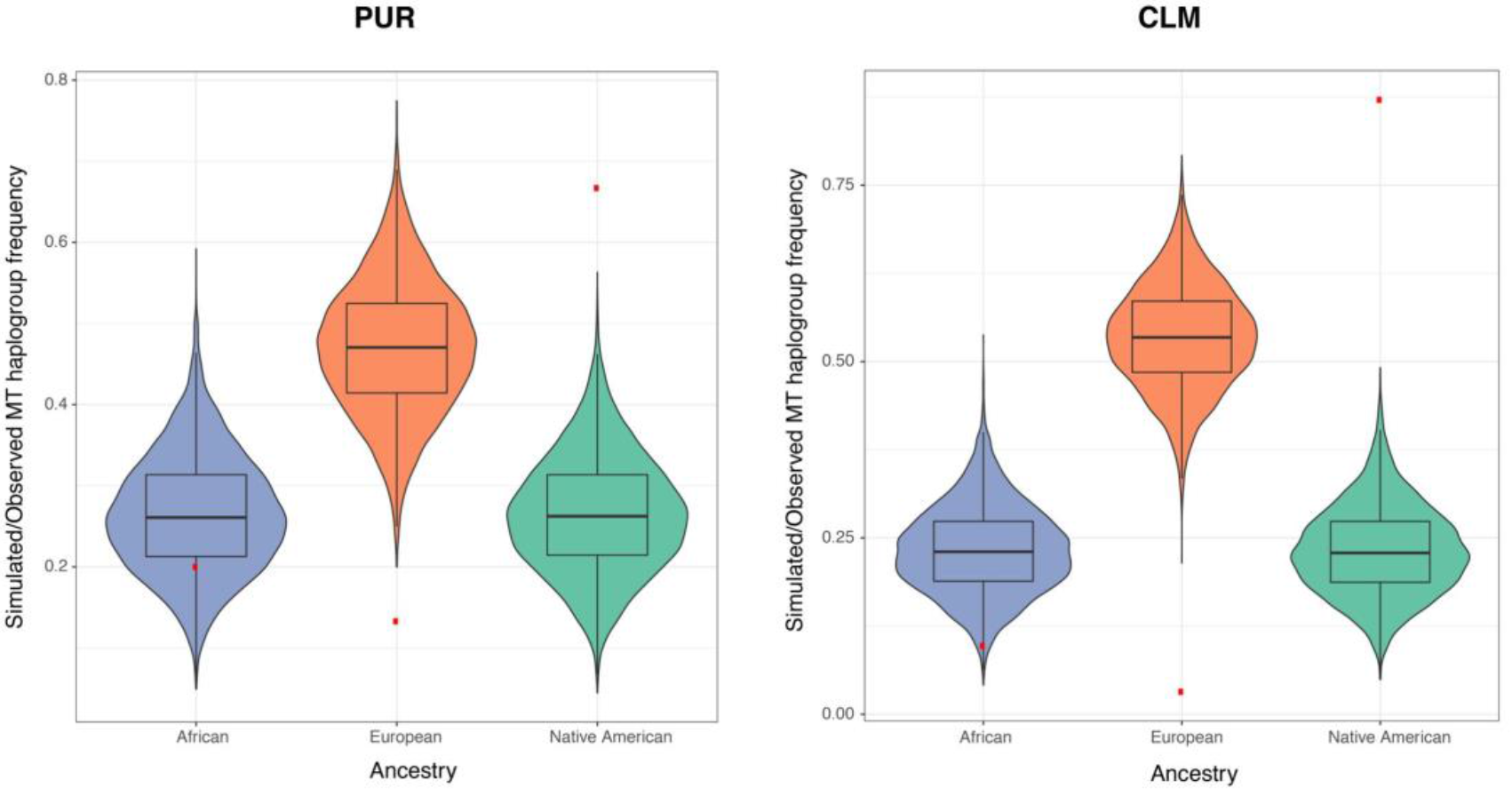
Results of simulations showing that drift since admixture is not sufficient to account for the increase in frequency of Native American mtDNA haplogroup, and concomitant decrease in European mtDNA haplogroup, in Colombians and Puerto Ricans. The violin plots show ancestry for 10,000 independent loci evolving neutrally since admixture. Thus, the median of the distribution represents the average ancestry contribution from each source population (African, European, and Native American), and the width of the distribution represents the amount of drift in ancestry since admixture. The red points indicate the observed frequency of mtDNA haplogroups.

## References

1. Wallace, D.C. (2007). Why do we still have a maternally inherited mitochondrial DNA? Insights from evolutionary medicine. Annu. Rev. Biochem. 76, 781–821.

2. Schatz, G. (2008). The protein import machinery of mitochondria. Protein Sci. 2, 141–146.

3. Quirós, P.M., Mottis, A., and Auwerx, J. (2016). Mitonuclear communication in homeostasis and stress. Nat. Rev. Mol. Cell Biol. 17, 213–226.

4. Pfanner, N., and Geissler, A. (2001). Versatility of the mitochondrial protein import machinery. Nat. Rev. Mol. Cell Biol. 2, 339–349.

5. Willett, C.S., and Burton, R.S. (2004). Evolution of Interacting Proteins in the Mitochondrial Electron Transport System in a Marine Copepod. Mol. Biol. Evol. 21, 443–453.

6. Osada, N., and Akashi, H. (2012). Mitochondrial-nuclear interactions and accelerated compensatory evolution: evidence from the primate cytochrome C oxidase complex. Mol. Biol. Evol. 29, 337–346.

7. Barreto, F.S., and Burton, R.S. (2013). Evidence for compensatory evolution of ribosomal proteins in response to rapid divergence of mitochondrial rRNA. Mol. Biol. Evol. 30, 310–314.

8. Burton, R.S., and Barreto, F.S. (2012). A disproportionate role for mtDNA in Dobzhansky-Muller incompatibilities? Mol. Ecol. 21, 4942–4957.

9. Rand, D.M., Haney, R.A., and Fry, A.J. (2004). Cytonuclear coevolution: the genomics of cooperation. Trends Ecol. Evol. 19, 645–653.

10. Wolff, J.N., Ladoukakis, E.D., Enríquez, J.A., and Dowling, D.K. (2014). Mitonuclear interactions: evolutionary consequences over multiple biological scales. Philos. Trans. R. Soc. Lond. B Biol. Sci. 369, 20130443.

11. Levin, L., Blumberg, A., Barshad, G., and Mishmar, D. (2014). Mito-nuclear co-evolution: the positive and negative sides of functional ancient mutations. Front. Genet. 5, 448.

12. Sackton, T.B., Haney, R.A., and Rand, D.M. (2003). Cytonuclear coadaptation in Drosophila: disruption of cytochrome c oxidase activity in backcross genotypes. Evolution 57, 2315–2325.

13. Mossman, J.A., Tross, J.G., Jourjine, N.A., Li, N., Wu, Z., and Rand, D.M. (2017). Mitonuclear Interactions Mediate Transcriptional Responses to Hypoxia in Drosophila. Mol. Biol. Evol. 34, 447–466.

14. Meiklejohn, C.D., Holmbeck, M.A., Siddiq, M.A., Abt, D.N., Rand, D.M., and Montooth, K.L. (2013). An Incompatibility between a mitochondrial tRNA and its nuclear-encoded tRNA synthetase compromises development and fitness in Drosophila. PLoS Genet. 9, e1003238.

15. James, A.C., and Ballard, J.W.O. (2003). Mitochondrial genotype affects fitness in Drosophila simulans. Genetics 164, 187–194.

16. Montooth, K.L., Meiklejohn, C.D., Abt, D.N., and Rand, D.M. (2010). Mitochondrial-nuclear epistasis affects fitness within species but does not contribute to fixed incompatibilities between species of Drosophila. Evolution 64, 3364–3379.

17. Dowling, D.K., Friberg, U., Hailer, F., and Arnqvist, G. (2007). Intergenomic epistasis for fitness: within-population interactions between cytoplasmic and nuclear genes in Drosophila melanogaster. Genetics 175, 235–244.

18. Hoekstra, L.A., Siddiq, M.A., and Montooth, K.L. (2013). Pleiotropic Effects of a Mitochondrial–Nuclear Incompatibility Depend upon the Accelerating Effect of Temperature in Drosophila. Genetics 195, 1129–1139.

19. Ellison, C.K., and Burton, R.S. (2006). Disruption of mitochondrial function in interpopulation hybrids of Tigriopus californicus. Evolution 60, 1382–1391.

20. Ellison, C.K., and Burton, R.S. (2008). Interpopulation hybrid breakdown maps to the mitochondrial genome. Evolution 62, 631–638.

21. Barreto, F.S., and Burton, R.S. (2013). Elevated oxidative damage is correlated with reduced fitness in interpopulation hybrids of a marine copepod. Proc. Biol. Sci. 280, 20131521.

22. Rawson, P.D., and Burton, R.S. (2002). Functional coadaptation between cytochrome c and cytochrome c oxidase within allopatric populations of a marine copepod. Proc. Natl. Acad. Sci. U. S. A. 99, 12955–12958.

23. Chou, J.-Y., and Leu, J.-Y. (2010). Speciation through cytonuclear incompatibility: insights from yeast and implications for higher eukaryotes. Bioessays 32, 401–411.

24. Lee, H.-Y., Chou, J.-Y., Cheong, L., Chang, N.-H., Yang, S.-Y., and Leu, J.-Y. (2008). Incompatibility of nuclear and mitochondrial genomes causes hybrid sterility between two yeast species. Cell 135, 1065–1073.

25. Chou, J.-Y., Hung, Y.-S., Lin, K.-H., Lee, H.-Y., and Leu, J.-Y. (2010). Multiple Molecular Mechanisms Cause Reproductive Isolation between Three Yeast Species. PLoS Biol. 8, e1000432.

26. Ellison, C.K., and Burton, R.S. (2010). Cytonuclear conflict in interpopulation hybrids: the role of RNA polymerase in mtDNA transcription and replication. J. Evol. Biol. 23, 528–538.

27. Morales, H.E., Pavlova, A., Amos, N., Major, R., Kilian, A., Greening, C., and Sunnucks, P. (2016). Mitochondrial-nuclear interactions maintain geographic separation of deeply diverged mitochondrial lineages in the face of nuclear gene flow.

28. Baris, T.Z., Wagner, D.N., Dayan, D.I., Du, X., Blier, P.U., Pichaud, N., Oleksiak, M.F., and Crawford, D.L. (2017). Evolved genetic and phenotypic differences due to mitochondrial-nuclear interactions. PLoS Genet. 13, e1006517.

29. Gershoni, M., Levin, L., Ovadia, O., Toiw, Y., Shani, N., Dadon, S., Barzilai, N., Bergman, A., Atzmon, G., Wainstein, J., et al. (2014). Disrupting Mitochondrial–Nuclear Coevolution Affects OXPHOS Complex I Integrity and Impacts Human Health. Genome Biol. Evol. 6, 2665–2680.

30. Sloan, D.B., Fields, P.D., and Havird, J.C. (2015). Mitonuclear linkage disequilibrium in human populations. Proc. Biol. Sci. 282,.

31. Rosenberg, N.A. (2002). Genetic Structure of Human Populations. Science 298, 2381–2385.

32. Cann, H.M. (2002). A Human Genome Diversity Cell Line Panel. Science 296, 261b–262.

33. Sharbrough, J., Havird, J.C., Noe, G.R., Warren, J.M., and Sloan, D.B. (2017). The Mitonuclear Dimension of Neanderthal and Denisovan Ancestry in Modern Human Genomes. Genome Biol. Evol. 9, 1567–1581.

34. Serre, D., Langaney, A., Chech, M., Teschler-Nicola, M., Paunovic, M., Mennecier, P., Hofreiter, M., Possnert, G., and Pääbo, S. (2004). No evidence of Neandertal mtDNA contribution to early modern humans. PLoS Biol. 2, E57.

35. Krings, M., Stone, A., Schmitz, R.W., Krainitzki, H., Stoneking, M., and Pääbo, S. (1997). Neandertal DNA sequences and the origin of modern humans. Cell 90, 19–30.

36. Darvasi, A., and Shifman, S. (2005). The beauty of admixture. Nat. Genet. 37, 118–119.

37. Shriner, D. (2013). Overview of admixture mapping. Curr. Protoc. Hum. Genet. Chapter 1, Unit 1.23.

38. Sans, M. (2000). Admixture studies in Latin America: from the 20th to the 21st century. Hum. Biol. 72, 155–177.

39. Mendizabal, I., Sandoval, K., Berniell-Lee, G., Calafell, F., Salas, A., Martínez-Fuentes, A., and Comas, D. (2008). Genetic origin, admixture, and asymmetry in maternal and paternal human lineages in Cuba. BMC Evol. Biol. 8, 213.

40. Sans, M., Weimer, T.A., Franco, M.H.L.P., Salzano, F.M., Bentancor, N., Alvarez, I., Bianchi, N.O., and Chakraborty, R. (2002). Unequal contributions of male and female gene pools from parental populations in the African descendants of the city of Melo, Uruguay. Am. J. Phys. Anthropol. 118, 33–44.

41. González-Andrade, F., Sánchez, D., González-Solórzano, J., Gascón, S., and Martínez-Jarreta, B. (2007). Sex-specific genetic admixture of Mestizos, Amerindian Kichwas, and Afro-Ecuadorans from Ecuador. Hum. Biol. 79, 51–77.

42. Dipierri, J.E., Alfaro, E., Martínez-Marignac, V.L., Bailliet, G., Bravi, C.M., Cejas, S., and Bianchi, N.O. (1998). Paternal directional mating in two Amerindian subpopulations located at different altitudes in northwestern Argentina. Hum. Biol. 70, 1001–1010.

43. Alves-Silva, J., da Silva Santos, M., Guimarães, P.E., Ferreira, A.C., Bandelt, H.J., Pena, S.D., and Prado, V.F. (2000). The ancestry of Brazilian mtDNA lineages. Am. J. Hum. Genet. 67, 444–461.

44. Moreno-Estrada, A., Gravel, S., Zakharia, F., McCauley, J.L., Byrnes, J.K., Gignoux, C.R., Ortiz-Tello, P.A., Martínez, R.J., Hedges, D.J., Morris, R.W., et al. (2013). Reconstructing the population genetic history of the Caribbean. PLoS Genet. 9, e1003925.

45. Homburger, J.R., Moreno-Estrada, A., Gignoux, C.R., Nelson, D., Sanchez, E., Ortiz-Tello, P., Pons-Estel, B.A., Acevedo-Vasquez, E., Miranda, P., Langefeld, C.D., et al. (2015). Genomic Insights into the Ancestry and Demographic History of South America. PLoS Genet. 11, e1005602.

46. Ruiz-Linares, A., Adhikari, K., Acuña-Alonzo, V., Quinto-Sanchez, M., Jaramillo, C., Arias, W., Fuentes, M., Pizarro, M., Everardo, P., de Avila, F., et al. (2014). Admixture in Latin America: geographic structure, phenotypic diversity and self-perception of ancestry based on 7,342 individuals. PLoS Genet. 10, e1004572.

47. Mörner, M. (1975). Race Mixture in the History of Latin America.

48. Malik, A.N., and Czajka, A. (2013). Is mitochondrial DNA content a potential biomarker of mitochondrial dysfunction? Mitochondrion 13, 481–492.

49. Yu, M. (2011). Generation, function and diagnostic value of mitochondrial DNA copy number alterations in human cancers. Life Sci. 89, 65–71.

50. Hu, L., Yao, X., and Shen, Y. (2016). Altered mitochondrial DNA copy number contributes to human cancer risk: evidence from an updated meta-analysis. Sci. Rep. 6, 35859.

51. Mengel-From, J., Thinggaard, M., Dalgård, C., Kyvik, K.O., Christensen, K., and Christiansen, L. (2014). Mitochondrial DNA copy number in peripheral blood cells declines with age and is associated with general health among elderly. Hum. Genet. 133, 1149–1159.

52. Joesch-Cohen, L.M., and Glusman, G. (2017). Differences between the genomes of lymphoblastoid cell lines and blood-derived samples. Adv. Genomics Genet. 7, 1–9.

53. Chakrabarty, S., D’Souza, R.R., Kabekkodu, S.P., Gopinath, P.M., Rossignol, R., and Satyamoorthy, K. (2014). Upregulation of TFAM and mitochondrial copy number in human lymphoblastoid cells. Mitochondrion 15, 52–58.

54. Nickles, D., Madireddy, L., Yang, S., Khankhanian, P., Lincoln, S., Hauser, S.L., Oksenberg, J.R., and Baranzini, S.E. (2012). In depth comparison of an individual’s DNA and its lymphoblastoid cell line using whole genome sequencing. BMC Genomics 13, 477.

55. Alexander, D.H., Novembre, J., and Lange, K. (2009). Fast model-based estimation of ancestry in unrelated individuals. Genome Res. 19, 1655–1664.

56. Mao, X., Bigham, A.W., Mei, R., Gutierrez, G., Weiss, K.M., Brutsaert, T.D., Leon-Velarde, F., Moore, L.G., Vargas, E., McKeigue, P.M., et al. (2007). A genomewide admixture mapping panel for Hispanic/Latino populations. Am. J. Hum. Genet. 80, 1171–1178.

57. Purcell, S., Neale, B., Todd-Brown, K., Thomas, L., Ferreira, M.A.R., Bender, D., Maller, J., Sklar, P., de Bakker, P.I.W., Daly, M.J., et al. (2007). PLINK: a tool set for whole-genome association and population-based linkage analyses. Am. J. Hum. Genet. 81, 559–575.

58. Chang, C.C., Chow, C.C., Tellier, L.C., Vattikuti, S., Purcell, S.M., and Lee, J.J. (2015). Second-generation PLINK: rising to the challenge of larger and richer datasets. Gigascience 4, 7.

59. Kloss-Brandstätter, A., Pacher, D., Schönherr, S., Weissensteiner, H., Binna, R., Specht, G., and Kronenberg, F. (2011). HaploGrep: a fast and reliable algorithm for automatic classification of mitochondrial DNA haplogroups. Hum. Mutat. 32, 25–32.

60. Maples, B.K., Gravel, S., Kenny, E.E., and Bustamante, C.D. (2013). RFMix: a discriminative modeling approach for rapid and robust local-ancestry inference. Am. J. Hum. Genet. 93, 278–288.

61. Martin, A.R., Gignoux, C.R., Walters, R.K., Wojcik, G.L., Neale, B.M., Gravel, S., Daly, M.J., Bustamante, C.D., and Kenny, E.E. (2017). Human Demographic History Impacts Genetic Risk Prediction across Diverse Populations. Am. J. Hum. Genet. 100, 635–649.

62. Bryc, K., Durand, E.Y., Michael Macpherson, J., Reich, D., and Mountain, J.L. (2015). The Genetic Ancestry of African Americans, Latinos, and European Americans across the United States. Am. J. Hum. Genet. 96, 37–53.

63. Lind, J.M., Hutcheson-Dilks, H.B., Williams, S.M., Moore, J.H., Essex, M., Ruiz-Pesini, E., Wallace, D.C., Tishkoff, S.A., O’Brien, S.J., and Smith, M.W. (2007). Elevated male European and female African contributions to the genomes of African American individuals. Hum. Genet. 120, 713–722.

64. Long, J.C. (1991). The genetic structure of admixed populations. Genetics 127, 417–428.

65. Pfaff, C.L., Parra, E.J., Bonilla, C., Hiester, K., McKeigue, P.M., Kamboh, M.I., Hutchinson, R.G., Ferrell, R.E., Boerwinkle, E., and Shriver, M.D. (2001). Population structure in admixed populations: effect of admixture dynamics on the pattern of linkage disequilibrium. Am. J. Hum. Genet. 68, 198–207.

66. Calvo, S.E., Clauser, K.R., and Mootha, V.K. (2015). MitoCarta2.0: an updated inventory of mammalian mitochondrial proteins. Nucleic Acids Res. 44, D1251–D1257.

67. Quinlan, A.R. (2014). BEDTools: The Swiss-Army Tool for Genome Feature Analysis. Curr. Protoc. Bioinformatics 47, 11.12.1–34.

68. 1000 Genomes Project Consortium, Auton, A., Brooks, L.D., Durbin, R.M., Garrison, E.P., Kang, H.M., Korbel, J.O., Marchini, J.L., McCarthy, S., McVean, G.A., et al. (2015). A global reference for human genetic variation. Nature 526, 68–74.

69. Rishishwar, L., and Jordan, I.K. (2017). Implications of human evolution and admixture for mitochondrial replacement therapy. BMC Genomics 18, 140.

70. Bailey, L.J., and Doherty, A.J. (2017). Mitochondrial DNA replication: a PrimPol perspective. Biochem. Soc. Trans. 45, 513–529.

71. Ciesielski, G.L., Oliveira, M.T., and Kaguni, L.S. (2016). Animal Mitochondrial DNA Replication. Enzymes 39, 255–292.

72. Jeon, J.-P., Shim, S.-M., Nam, H.-Y., Baik, S.-Y., Kim, J.-W., and Han, B.-G. (2007). Copy number increase of 1p36.33 and mitochondrial genome amplification in Epstein–Barr virus-transformed lymphoblastoid cell lines. Cancer Genet. Cytogenet. 173, 122–130.

73. Bhatia, G., Patterson, N., Sankararaman, S., and Price, A.L. (2013). Estimating and interpreting FST: the impact of rare variants. Genome Res. 23, 1514–1521.

74. Pagliarini, D.J., Calvo, S.E., Chang, B., Sheth, S.A., Vafai, S.B., Ong, S.-E., Walford, G.A., Sugiana, C., Boneh, A., Chen, W.K., et al. (2008). A mitochondrial protein compendium elucidates complex I disease biology. Cell 134, 112–123.

75. Holt, I.J., and Reyes, A. (2012). Human mitochondrial DNA replication. Cold Spring Harb. Perspect. Biol. 4,.

76. Suissa, S., Wang, Z., Poole, J., Wittkopp, S., Feder, J., Shutt, T.E., Wallace, D.C., Shadel, G.S., and Mishmar, D. (2009). Ancient mtDNA genetic variants modulate mtDNA transcription and replication. PLoS Genet. 5, e1000474.

77. Trounce, I., Neill, S., and Wallace, D.C. (1994). Cytoplasmic transfer of the mtDNA nt 8993 T-->G (ATP6) point mutation associated with Leigh syndrome into mtDNA-less cells demonstrates cosegregation with a decrease in state III respiration and ADP/O ratio. Proc. Natl. Acad. Sci. U. S. A. 91, 8334–8338.

78. Eyre-Walker, A. (2017). Mitochondrial Replacement Therapy: Are Mito-nuclear Interactions Likely To Be a Problem? Genetics 205, 1365–1372.

79. Lopez-Lluch, G., Hunt, N., Jones, B., Zhu, M., Jamieson, H., Hilmer, S., Cascajo, M.V., Allard, J., Ingram, D.K., Navas, P., et al. (2006). Calorie restriction induces mitochondrial biogenesis and bioenergetic efficiency. Proceedings of the National Academy of Sciences 103, 1768–1773.

80. Lee, H.-C., and Wei, Y.-H. (2005). Mitochondrial biogenesis and mitochondrial DNA maintenance of mammalian cells under oxidative stress. Int. J. Biochem. Cell Biol. 37, 822–834.

81. Wu, Z., Puigserver, P., Andersson, U., Zhang, C., Adelmant, G., Mootha, V., Troy, A., Cinti, S., Lowell, B., Scarpulla, R.C., et al. (1999). Mechanisms controlling mitochondrial biogenesis and respiration through the thermogenic coactivator PGC-1. Cell 98, 115–124.

82. Florez, J.C., Price, A.L., Campbell, D., Riba, L., Parra, M.V., Yu, F., Duque, C., Saxena, R., Gallego, N., Tello-Ruiz, M., et al. (2009). Strong association of socioeconomic status with genetic ancestry in Latinos: implications for admixture studies of type 2 diabetes. Diabetologia 52, 1528–1536.

83. Mishmar, D., Ruiz-Pesini, E., Golik, P., Macaulay, V., Clark, A.G., Hosseini, S., Brandon, M., Easley, K., Chen, E., Brown, M.D., et al. (2003). Natural selection shaped regional mtDNA variation in humans. Proc. Natl. Acad. Sci. U. S. A. 100, 171–176.

84. Balloux, F., Handley, L.-J.L., Jombart, T., Liu, H., and Manica, A. (2009). Climate shaped the worldwide distribution of human mitochondrial DNA sequence variation. Proc. Biol. Sci. 276, 3447–3455.

85. Jobling, M., Hurles, M., and Tyler-Smith, C. (2013). Human Evolutionary Genetics: Origins, Peoples & Disease (Garland Science).

86. Brown, G.R., Laland, K.N., and Mulder, M.B. (2009). Bateman’s principles and human sex roles. Trends Ecol. Evol. 24, 297–304.

87. Betzig, L. (2012). Means, variances, and ranges in reproductive success: comparative evidence. Evol. Hum. Behav. 33, 309–317.

88. Sudlow, C., Gallacher, J., Allen, N., Beral, V., Burton, P., Danesh, J., Downey, P., Elliott, P., Green, J., Landray, M., et al. (2015). UK biobank: an open access resource for identifying the causes of a wide range of complex diseases of middle and old age. PLoS Med. 12, e1001779.

89. Reinhardt, K., Dowling, D.K., and Morrow, E.H. (2013). Medicine. Mitochondrial replacement, evolution, and the clinic. Science 341, 1345–1346.

90. Wolf, D.P., Mitalipov, N., and Mitalipov, S. (2015). Mitochondrial replacement therapy in reproductive medicine. Trends Mol. Med. 21, 68–76.

91. Gemmell, N., and Wolff, J.N. (2015). Mitochondrial replacement therapy: Cautiously replace the master manipulator. Bioessays 37, 584–585.

92. Kenney, M.C., Chwa, M., Atilano, S.R., Falatoonzadeh, P., Ramirez, C., Malik, D., Tarek, M., Del Carpio, J.C., Nesburn, A.B., Boyer, D.S., et al. (2014). Molecular and bioenergetic differences between cells with African versus European inherited mitochondrial DNA haplogroups: implications for population susceptibility to diseases. Biochim. Biophys. Acta 1842, 208–219.

93. Ballinger, S.W. (2013). Beyond retrograde and anterograde signalling: mitochondrial-nuclear interactions as a means for evolutionary adaptation and contemporary disease susceptibility. Biochem. Soc. Trans. 41, 111–117.

